# Gauge fixing for sequence-function relationships

**DOI:** 10.1101/2024.05.12.593772

**Authors:** Anna Posfai, Juannan Zhou, David M. McCandlish, Justin B. Kinney

**Author notes:** Correspondence (DMM), (JBK). Please provide details of author contributions here. Please declare any competing interests here.

## Abstract

Quantitative models of sequence-function relationships are ubiquitous in computational biology, e.g., for modeling the DNA binding of transcription factors or the fitness landscapes of proteins. Interpreting these models, however, is complicated by the fact that the values of model parameters can often be changed without affecting model predictions. Before the values of model parameters can be meaningfully interpreted, one must remove these degrees of freedom (called “gauge freedoms” in physics) by imposing additional constraints (a process called “fixing the gauge”). However, strategies for fixing the gauge of sequence-function relationships have received little attention. Here we derive an analytically tractable family of gauges for a large class of sequence-function relationships. These gauges are derived in the context of models with all-order interactions, but an important subset of these gauges can be applied to diverse types of models, including additive models, pairwise-interaction models, and models with higher-order interactions. Many commonly used gauges are special cases of gauges within this family. We demonstrate the utility of this family of gauges by showing how different choices of gauge can be used both to explore complex activity landscapes and to reveal simplified models that are approximately correct within localized regions of sequence space. The results provide practical gauge-fixing strategies and demonstrate the utility of gauge-fixing for model exploration and interpretation.

**Significance Statement:** Computational biology relies heavily on mathematical models that predict biological activities from DNA, RNA, or protein sequences. Interpreting the parameters of these models, however, remains difficult. Here we address a core challenge for model interpretation-the presence of ‘gauge freedoms’, i.e., ways of changing model parameters without affecting model predictions. The results unify commonly used methods for eliminating gauge freedoms and show how these methods can be used to simplify complex models in localized regions of sequence space. This work thus overcomes a major obstacle in the interpretation of quantitative sequence-function relationships.

## Introduction

One of the central challenges of biology is to understand how functionally relevant information is encoded within the sequences of DNA, RNA, and proteins. Unlike the genetic code, most sequence-function relationships are quantitative in nature, and understanding them requires finding mathematical functions that, upon being fed unannotated sequences, return values that quantify sequence activity (1). Multiplex assays of variant effects (MAVEs), functional genomics methods, and other high-throughput techniques are rapidly increasing the ease with which sequence-function relationships can be experimentally studied. And while quantitative modeling efforts based on these high-throughput data are becoming increasingly successful, in that they yield models with ever-increasing predictive ability, major open questions remain about how to interpret both the parameters (2–12) and the predictions (13–17) of the resulting models. One major open question is how to deal with the presence of gauge freedoms.

Gauge freedoms are directions in parameter space along which changes in model parameters have no effect on model predictions (18). Not only can the values of model parameters along gauge freedoms not be determined from data, differences in parameters along gauge freedoms have no biological meaning even in principle. Many commonly used models of sequence-function relationships exhibit numerous gauge freedoms (19– 35), and interpreting the parameters of these models requires imposing additional constraints on parameter values, a process called “fixing the gauge”.

The gauge freedoms of sequence-function relationships are currently most completely understood in the context of additive models [commonly used to describe transcription factor binding to DNA (19, 22, 35)] and pairwise-interaction models [commonly used to describe proteins (20, 21, 23–34)]. Recently, some gauge-fixing strategies have been described for all-order interaction models, again in the context of protein sequence-function relationships (30, 31, 34). However, a unified gauge-fixing strategy applicable to diverse models of sequence-function relationships has yet to be developed.

Here we provide a general treatment of the gauge fixing problem for sequence-function relationships, focusing on the important case where the set of gauge-fixed parameters form a vector space, thus ensuring that differences between vectors of gauge-fixed parameter values are directly interpretable. We first demonstrate the relationship between these linear gauges and *L*_2_ regularization on parameter vectors, and then derive a mathematically tractable family of gauges for the all-order interaction model. Importantly, a subset of these gauges–the “hierarchical gauges”–can be applied to diverse lower-order models (including additive models, pairwise-interaction models, and higher-order interaction models) and include as special cases two types of gauges that are commonly used in practice [“zero-sum gauges” (23, 28) and “wild-type gauges” (9, 23, 33)]. We then illustrate the properties of this family of gauges by analyzing two example sequence-function relationships: a simulated all-order interaction landscape on short binary sequences, and an empirical pairwise-interaction landscape for the B1 domain of protein G (GB1). The GB1 analysis, in particular, shows how different hierarchical gauges can be used to explore, simplify, and interpret complex functional landscapes. A companion paper (36) further explores the mathematical origins of gauge freedoms in models of sequence-function relationships, and shows how gauge freedoms arise as a consequence of the symmetries of sequence space.

## Results

### Preliminaries and background

In this section we review how gauge freedoms arise in commonly used models of sequence-function relationships, as well as strategies commonly used to fix the gauge. In doing so, we establish notation and concepts that are used in subsequent sections, as well as in our companion paper (36).

#### Linear models

We define quantitative models of sequence-function relationships as follows. Let *A* denote an alphabet comprising *α* distinct characters (written *c*_1_, …, *c*_*α*_), let 𝒮 denote the set of sequences of length *L* built from these characters, and let *N* = *α*^*L*^ denote the number of sequences in 𝒮. A quantitative model of a sequence-function relationship (henceforth “model”) is a function 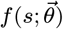 that maps each sequence *s* in 𝒮 to a real number. The vector 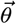 represents the parameters on which this function depends and is assumed to comprise *M* real numbers. *s*_*l*_ denotes the character at position *l* of sequence *s*. We use *l, l*^*′*^, etc. to index positions (ranging from 1 to *L*) in a sequence and *c, c*^*′*^, etc. to index characters in 𝒜.

A linear model is a model that is a linear function of 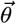. Linear models have the form

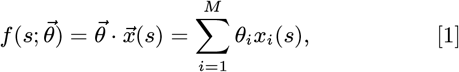

where 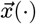 is a vector of *M* distinct sequence features and each sequence feature *x*_*i*_(·) is a function that maps sequences to the real numbers. We refer to the space ℝ^*M*^ in which 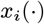 lives as feature space, and the specific vector 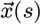 as the embedding of sequence *s* in feature space. We use *S* to denote the vector space spanned by the set of embeddings 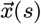 for all sequences *s* in 𝒮.

#### One-hot models

One-hot models are linear models based on sequence features that indicate the presence or absence of specific characters at specific positions within a sequence (1). Such models play a central role in scientific reasoning concerning sequence-function relationships because their parameters can be interpreted as quantitative contributions to the measured function due to the presence of specific biochemical entities (e.g. nucleotides or amino acids) in specific positions in the sequence. These one-hot models include additive models, pairwise-interaction models, all-order interaction models, and more. Additive models have the form

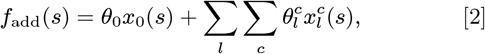

where *x*_0_(*s*) is the constant feature (equal to one for every sequence *s*) and 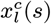 is an additive feature (equal to one if sequence *s* has character *c* at position *l* and equal to zero otherwise). Pairwise interaction models have the form

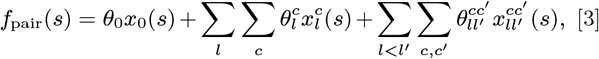

where 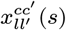 is a pairwise feature (equal to one if *s* has character *c* at position *l* and character *c*^*′*^ at position *l*^*′*^, and equal to zero otherwise). All-order interaction models include interactions of all orders, and are written

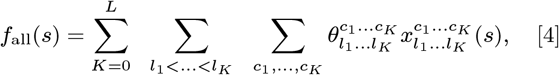

where 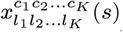 is a *K*-order feature (equal to one if *s* has character *c*_*k*_ at position *l*_*k*_ for all *k*, and equal to zero otherwise; *K* = 0 corresponds to the constant feature).

#### Gauge freedoms

Gauge freedoms are transformations of model parameters that leave all model predictions unchanged. The gauge freedoms of a general sequence-function relationship *f*(*·, ·*) are vectors 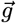 in ℝ^*M*^ that satisfy

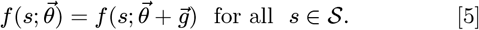

For linear models, gauge freedoms 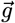 satisfy

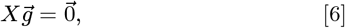

where *X* is the *N × M* design matrix having rows 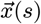 or *s* ∈ 𝒮. In linear models, gauge freedoms thus arise when sequence features (i.e., the columns of *X*) are not linearly independent. In such cases, the space *S* spanned by sequence embeddings is a proper subspace of ℝ^*M*^, so is the space *G* of gauge freedoms, and *G* is orthogonal to *S*.

Each linear relation between multiple columns of *X* yields a gauge freedom. For example, additive models have *L* gauge freedoms arising from the *L* linear relations,

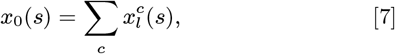

for all positions *l*. Pairwise models have *L* gauge freedoms arising from the *L* additive model linear relations in Eq. (7), and 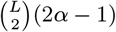 additional gauge freedoms arising from the linear relations

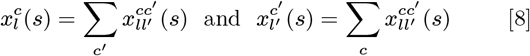

for all characters *c, c*^*′*^ and all positions *l* and *l*^*′*^, with *l < l*^*′*^ (see SI Sec. 2 for details). More generally, the gauge freedoms of one-hot models arise from the fact that summing any *K*-order feature 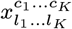 over all characters *c*_*k*_ at any chosen position *l*_*k*_ yields a feature of order *K* − 1.

#### Parameter values depend on choice of gauge

Gauge freedoms pose problems for the interpretation of model parameters because different choices of model parameters can give the exact same predictions when they are present. Thus, unless constraints are placed on the values of allowable parameters, individual parameters will have little biological meaning when viewed in isolation. To interpret model parameters, one therefore needs to adopt constraints that eliminate gauge freedoms and, as a result, make the values of model parameters unique. These constraints are called the “gauge” in which parameters are expressed, and this process of choosing constraints is called “fixing the gauge”. There are many different gauge-fixing strategies. For example, Fig. 1 shows an additive model of the DNA binding energy of CRP [an important transcription factor in *Escherichia coli* (37)] expressed in three different choices of gauge.

**Fig. 1.**
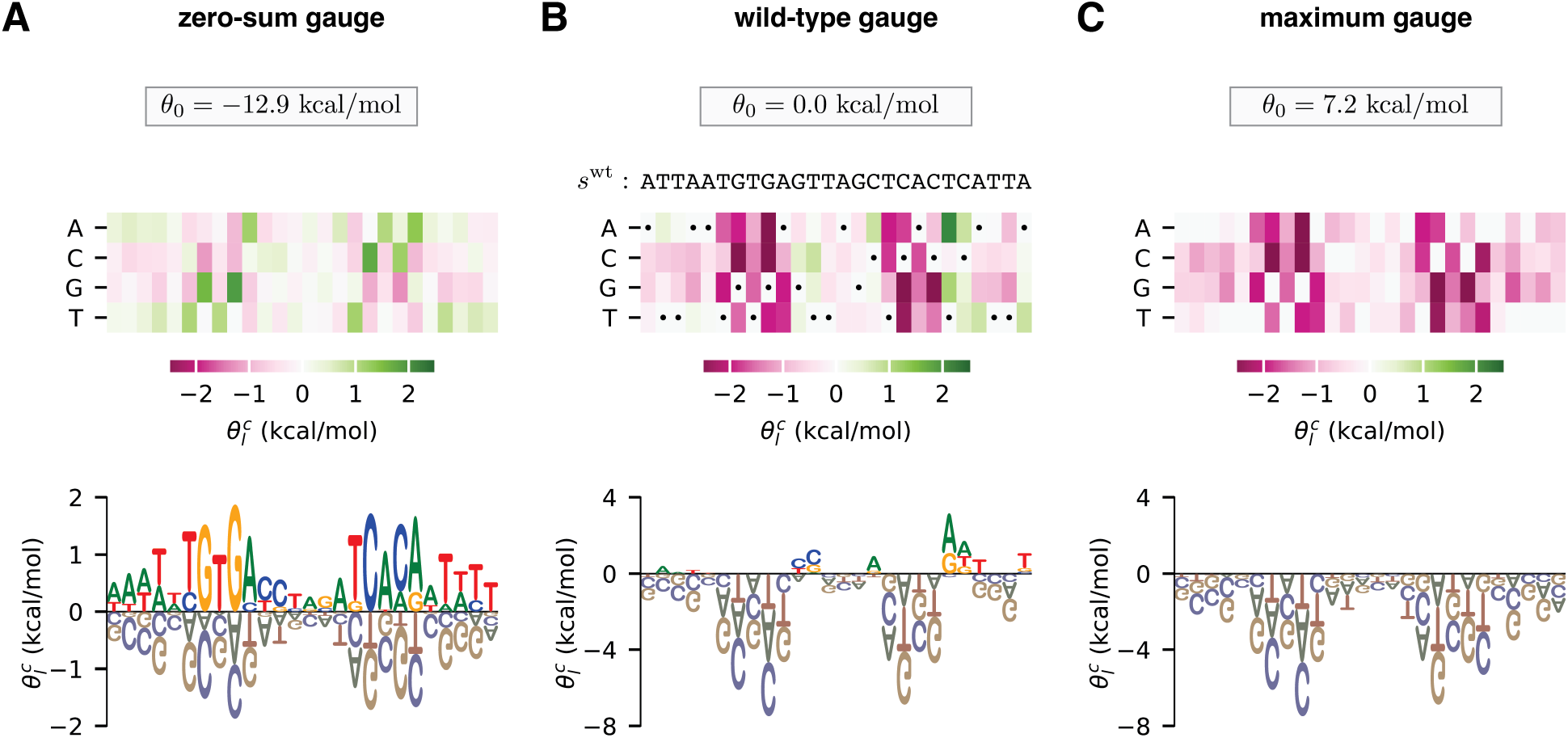
Choice of gauge impacts model parameters. (A-C) Parameters, expressed in three different gauges, for an additive model describing the (negative) binding energy of the *E. coli* transcription factor CRP to DNA. Model parameters are from (56). In each panel, additive parameters, 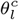, are shown using both (top) a heat map and (bottom) a sequence logo (57). The value of the constant parameter, *θ*_0_, is also shown. (A) The zero-sum gauge, in which the additive parameters at each position sum to zero. (B) The wild-type gauge, in which the additive parameters at each position quantify activity differences with respect to a wild-type sequence, *s*^wt^. The wild-type sequence used here (indicated by dots on the heat map) is the CRP binding site present at the *E. coli* lac promoter. (C) The maximum gauge, in which the additive parameters at each position quantify differences with respect to the optimal character at that position.

Fig. 1A shows parameters expressed in the “zero-sum gauge” (23, 28) [also called the “Ising gauge” (28), or the “hierarchical gauge” (9)]. In the zero-sum gauge, the constant parameter is the mean sequence activity and the additive parameters quantify deviations from this mean activity. The name of the gauge comes from the fact that the additive parameters at each position sum to zero. The zero-sum gauge is commonly used in additive models of protein-DNA binding (35, 38–43). As we will see, zero-sum gauges are readily defined for models with pairwise and higher-order interactions as well.

Fig. 1B shows parameters expressed in the “wild-type gauge” (9, 23, 33) [also called the “lattice-gas gauge” (28), or the “mismatch gauge” (35)]. In the wild-type gauge, the constant parameter is equal to the activity of a chosen wild-type sequence (denoted *s*^*wt*^), and additive parameters are the changes in activity that result from mutations away from the wild-type sequence. The wild-type gauge is commonly used to visualize the results of mutational scanning experiments on proteins (44–48) or on long DNA regulatory sequences (49–54). As we will see, wild-type gauges are also readily defined for models with pairwise and higher-order interactions.

Fig. 1C shows parameters expressed in what we call the “maximum gauge”. In the maximum gauge, the constant parameter is equal to the activity of the highest-activity sequence, and additive parameters are the changes in activity that result from mutations away from the highest-activity sequence. The maximum gauge is less common in the literature than the zero-sum and the wild-type gauge, but has been used in multiple publications (55, 56).

#### Gauge spaces

We now turn our attention to strategies for fixing the gauge. For every parameter vector 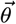 in ℝ^*M*^, there is a corresponding “gauge orbit” defined by the set of vectors that can be obtained from 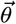 by adding a vector 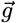 in the space of gauge freedoms *G*. We remove the gauge freedoms of a model (a process called “fixing the gauge”) by restricting valid parameter vectors to a specified “gauge space” Θ, a subset of ℝ^*M*^ that intersects the gauge orbit of each possible parameter vector 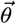 at exactly one point. That one point, denoted by 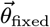, is called the “gauge-fixed” value of 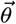.

For any model of a sequence-function relationship with gauge freedoms, there are an infinite number of possible choices for the gauge space Θ. Fig. 2 illustrates the three gauge spaces corresponding to the three different gauges (zero-sum, wild-type, and maximum) used in Fig. 1. In the zero-sum gauge (Fig. 2A), the α additive parameters at each position are restricted to a linear subspace of dimension α − 1 in which the sum of the parameters is zero. In the wild-type gauge (Fig. 2B), the additive parameters at each position are restricted to a linear subspace in which the parameters that contribute to the activity of the wild-type sequence are zero. In the maximum gauge (Fig. 2C), the additive parameters at each position are restricted to a nonlinear subspace in which all parameters are less than or equal to zero and, at every point in the subspace, at least one parameter is equal to zero.

**Fig. 2.**
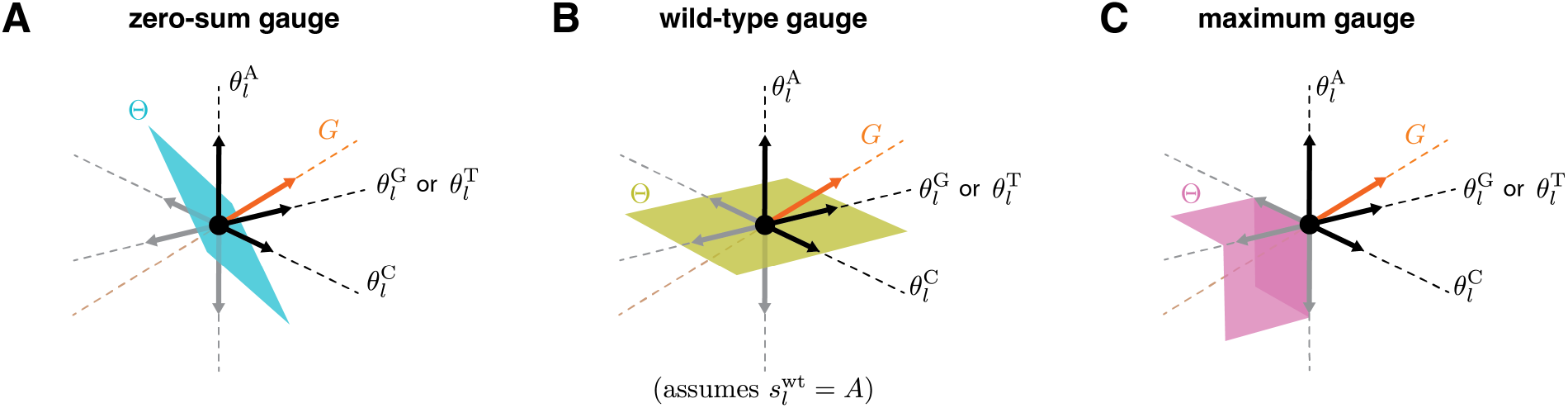
Geometry of gauge spaces for additive one-hot models. (A-C) Geometric representation of the gauge space Θ to which the additive parameters at each position *l* are restricted in the corresponding panel of Fig. 1. Each of the four sequence features (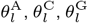, and 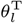) corresponds to a different axis. Note that the two axes for 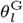 and 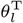 are shown as one axis to enable 3D visualization. Black and gray arrows respectively denote unit vectors pointing in the positive and negative directions along each axis. *G* indicates the space of gauge transformations.

#### Linear gauges

Here and throughout the rest of this paper we focus on linear gauges, i.e., choices of Θ that are linear sub-spaces of feature space (as in Fig. 2A,B). Linear gauges are the most mathematically tractable family of gauges. Linear gauges also have the attractive property that the difference between any two parameter vectors in Θ is also in Θ. This property makes the comparison of models within the same gauge straight-forward.

Parameters can be fixed to any chosen linear gauge via a corresponding linear projection. Formally, for any linear gauge Θ there exists an *M ×M* projection matrix *P* that projects any vector 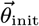 along the gauge space *G* to an equivalent vector 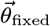 that lies in Θ, i.e.

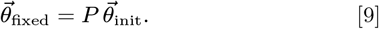

See SI Sec. 3 for a proof. We emphasize that *P* depends on the choice of Θ, and that *P* is an orthogonal projection only for the specific choice Θ = *S*.

Parameters can also be gauge-fixed through a process of constrained optimization. Let Λ be any positive-definite *M × M* matrix, and let 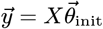 be the *N* -dimensional vector of model predictions on all sequences. Then Λ specifies a unique gauge-fixed set of parameters that preserves 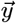 via

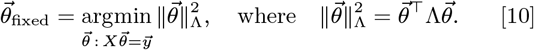

The resulting gauge space comprises the set of vectors that minimize the Λ-norm in each gauge orbit. The corresponding projection matrix is

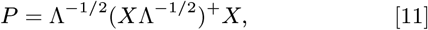

where ‘+’ indicates the Moore-Penrose pseudoinverse. See SI Sec. 3 for a proof. In what follows, the connection between the penalization matrix Λ and the projection matrix *P* will be used to help interpret the constraints imposed by the gauge space Θ.

One consequence of Eq. (10) is that parameter inference carried out using a positive-definite *L*_2_ regularizer Λ on model parameters will result in gauge-fixed model parameters in the specific linear gauge determined by Λ (see SI Sec. 3). While it might then seem that *L*2 regularizing parameter values during inference solves the gauge fixing problem, it is important to understand that regularizing during model inference will also change model predictions, whereas gauge-fixing proper influences only the model parameters while keeping the model predictions fixed. In addition, we show in SI Sec. 3 that, for any desired positive-definite regularizer on model predictions and choice of linear gauge Θ, we can construct a positive-definite penalization matrix for model parameters Λ that imposes the desired regularization on model predictions and yields inferred parameters in the desired gauge. Thus while *L*2 regularization during parameter inference can simultaneously fix the gauge and regularize model predictions, the regularization imposed on model predictions does not constrain the choice of gauge.

### Unified approach to gauge fixing

We now derive strategies for fixing the gauge of the all-order interaction model. We first introduce a geometric formulation of the all-order interaction model embedding. We then construct a parametric family of gauges for the all-order interaction model, and derive formulas for the corresponding projection and penalizing matrices. Next, we highlight specific gauges of interest in this parametric family. We focus in particular on the “hierarchical gauges,” which can be applied to a variety of commonly used models in addition to the all-order interaction model. The results provide explicit gauge-fixing formulae that can be applied to diverse quantitative models of sequence-function relationships.

#### All-order interaction models

To aid in our discussion of the all-order interaction model [Eq. (4)], we define an augmented alphabet 𝒜^*′*^ = {*, *c*_1_, …, *c*_*α*_}, where *c*_1_, …, *c*_*α*_ are the characters in 𝒜 and * is a wild-card character that is interpreted as matching any character in 𝒜. Let 𝒮^*′*^ denote the set of sequences of length *L* comprising characters from 𝒜^*′*^. For each augmented sequence *s*^*′*^ ∈ 𝒮^*′*^, we define the sequence feature *x*_*s*_^*′*^ (*s*) to be 1 if a sequence *s* matches the pattern described by *s*^*′*^ and to be 0 otherwise. In this way, each augmented sequence *s*^*′*^ serves as a regular expression against which bona fide sequences are compared.

Assigning one parameter *θ*_*s*_^*′*^ to each of the *M* = (α + 1)^*L*^ augmented sequences *s*^*′*^, the all-order interaction model can be expressed compactly as

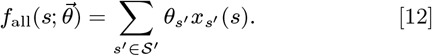

In this notation, the constant parameter θ_0_is written θ_*\*···\**_, each additive parameter 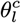 is written θ_*\*···c···\**_, each pairwise interaction parameter 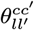 is written θ_*\*···c···c*_^*′*^_*···\**_, and so on. (Here *c* occurs at position *l, c*^*′*^ occurs at position *l*^*′*^, and · · · denotes a run of * characters). We thus see that augmented sequences provide a convenient way to index the features and parameters of the all-order interaction model.

Next we observe that *x*_*s*_^*′*^ can be expressed in a form that factorizes across positions. For each position *l*, we define 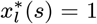 for all sequences *s* and take 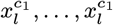 to be the standard one-hot sequence features. *x*_*s*_^*′*^ can then be written in the factorized form,

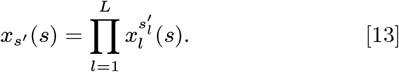

From this it is seen that the embedding for the all-order interaction model, 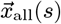, can be formulated geometrically as a tensor product:

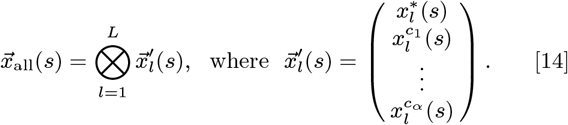

See SI Sec. 4 for details.

#### Parametric family of gauges

We now define a useful parametric family of gauges for the all-order interaction model. Each gauge in this family is defined by two parameters, λ and *p*. λ is a non-negative real number that governs how much higher-order versus lower-order sequence features are penalized [in the sense of Eq. (10)]. *p* is a probability distribution on sequence space that governs how strongly the specific characters at each position are penalized. This distribution is assumed to have the form

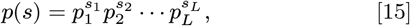

where 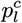 denotes the probability of character *c* at position *l*. As we show below, choosing appropriate values for *λ* and *p* recovers the most commonly used linear gauges, including the zero-sum gauge, the wild-type gauge, and more.

Gauges in the parametric family have analytically tractable projection matrices because they can be expressed as tensor products of single-position gauge spaces. Let 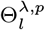 be the α-dimensional subspace of ℝ^α+1^ defined by

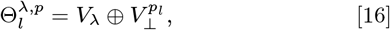

where *V*_*λ*_ (a 1-dimensional subspace) and 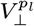 [an (α − 1)-dimensional subspace] are defined by

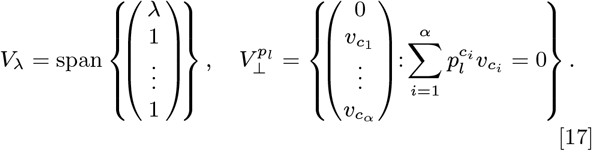

The full parametric gauge, denoted by Θ^*λ*,*p*^, is defined to be the tensor product of these single-position gauges:

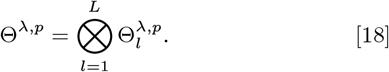

As detailed in SI Sec. 5, the corresponding projection matrix *P*^*λ*,*p*^ is found to have elements given by

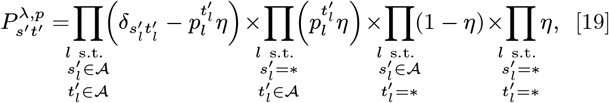

where η = λ/(1+λ) and where the augmented sequences *s*^*′*^ and *t*^*′*^ index rows and columns. We thus obtain an explicit formula for the projection matrix needed to project any parameter vector into any gauge in the parametric family.

Gauges in the parametric family also have penalizing matrices of a simple diagonal form. Specifically, if 0 < λ < ∞ and *p*(*s*^*′*^) > 0 everywhere, Eq. (10) is satisfied by the penalization matrix Λ having elements

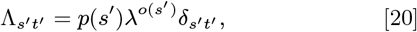

where *o*(*s*^*′*^) denotes the order of interaction described by *s*^*′*^ (i.e., the number of non-star characters in *s*^*′*^) and *p*(*s*^*′*^) is defined as in Eq. (15) but with 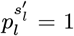 when 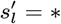. See SI Sec. 5 for a proof. Note that, although Eq. (20) does not hold when λ = 0, λ = ∞, or any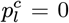, one can interpret Θ^*λ*,*p*^ [which is well-defined in Eq. (18) and Eq. (19)] as arising from Eq. (10) under a limiting series of penalizing matrices.

#### Trivial gauge

Choosing λ = 0 yields what we call the “trivial gauge”. In the trivial gauge, *θ*_*s*_^*′*^ = 0 if *s*^*′*^ contains one or more star characters (by Eq. (19)), and so the only nonzero parameters correspond to interactions of order *L*. As a result,

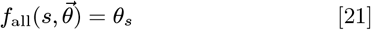

for every sequence *s* ∈ 𝒮. Note in particular that the trivial gauge is unaffected by *p*. Thus, the trivial gauge essentially represents sequence-function relationships as catalogs of activity values, one value for every sequence. See SI Sec. 6 for details.

#### Euclidean gauge

Choosing λ = α and choosing *p* to be the uniform distribution recovers what we call the “Euclidean gauge”. In the Euclidean gauge, the penalizing norm in Eq. (10) is the standard euclidean norm, i.e.

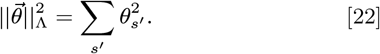

It is readily seen that the euclidean gauge is orthogonal to the space of gauge freedoms *G* and therefore equal to the embedding space *S*. It is also readily seen that parameter inference using standard *L*_2_ regularization (i.e. choosing Λ to be a positive multiple of the identity matrix) will yield parameters in the Euclidean gauge. See SI Sec. 6 for details.

#### Equitable gauge

Choosing λ = 1 and letting *p* vary recovers what we call the “equitable gauge”. In the equitable gauge, the penalizing norm is

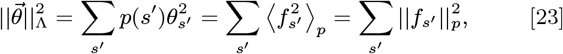

where *f*_*s*_^*′*^ = θ_*s*_^*′*^ *x*_*s*_^*′*^ denotes the contribution to the activity landscape corresponding to the sequence feature *s*^*′*^, ⟨·⟩_*p*_ denotes an average over sequences drawn from *p*, and 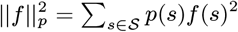 is the squared norm of a function *f* on sequence space with respect to *p*. The equitable gauge thus penalizes each parameter θ_*s*_^*′*^ in proportion to the fraction of sequences that parameter applies to. Equivalently, the equitable gauge can be thought of as minimizing the sum of the squared norms of the landscape contributions 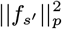 rather than the squared norm of the parameter values themselves. Unlike the euclidean gauge, the equitable gauge accounts for the fact that different model parameters can affect vastly different numbers of sequences and can thereby have vastly different impacts on the activity landscape. See SI Sec. 6 for details.

#### Hierarchical gauge

Choosing *p* freely and letting λ → ∞ yields what we call the “hierarchical gauge”. When expressed in the hierarchical gauge, model parameters obey the marginalization property,

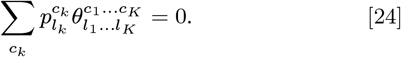

This marginalization property has important consequences that we now summarize. See SI Sec. 7 for proofs of these results.

A first consequence of Eq. (24) is that, when parameters are expressed in the hierarchical gauge, the mean activity among sequences matched by an augmented sequence *s*^*′*^ can be expressed as a simple sum of parameters. For example,

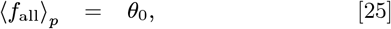

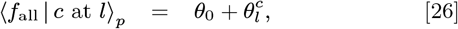

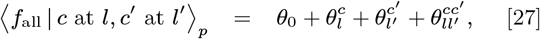

and so on. Consequently, the parameters themselves can also be expressed in terms of differences of these average values. For instance, 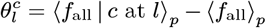. Because *p* factorizes by position, conditioning on having particular characters in a subset of positions is equivalent to the probability distribution produced by drawing sequences from *p* and then fixing those positions in the drawn sequences to those specific characters. Thus, 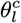 can also be interpreted as the average effect of mutating position *l* to character *c* when sequences are drawn from Similarly, 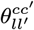 is the average effect of fixing positions *l* to *c* and *l*^*′*^ to *c*^*′*^ when drawing from *p* beyond what would be expected based on the effects of changing *l* to *c* and *l*^*′*^ to *c*^*′*^ individually (i.e. epistasis), and higher-order coefficients have a similar interpretation. The hierarchical gauge thus provides an ANOVA-like decomposition of activity landscapes.

A second consequence of Eq. (24) is that the activity landscape, when expressed in the hierarchical gauge, naturally decomposes into mutually orthogonal components. Let σ denote a set comprising all augmented sequences that have the same pattern of star and non-star positions, and let 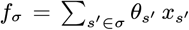 be the corresponding component of *f*_*all*_. These landscape components are *p*-orthogonal when expressed in the hierarchical gauge:

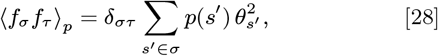

where σ and τ represent any two such sets of augmented sequences. One implication of this orthogonality relation is that the variance of the landscape (with respect to *p*) is the sum of contributions from interactions of different orders:

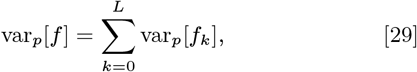

where *f*_*k*_ denotes the sum of *k*-order terms that contribute to *f*_*all*_. Another implication is that the hierarchical gauge minimizes the variance attributable to different orders of interaction in a hierarchical manner: higher-order terms are prioritized for variance minimization over lower-order terms, and within a given order parameters are penalized in proportion to the fraction of sequences they apply to.

A third consequence of Eq. (24) is that hierarchical gauges preserve the form of a large class of one-hot models that are equivalent to all-order interaction models with certain parameters fixed at zero (specifically, these models satisfy the condition that if a parameter for a sequence feature is fixed at zero, all higher-order sequence features contained within that sequence feature also have their parameters fixed at zero). These models, which we call the “hierarchical models,” include all-order interaction models in which the parameters above a specified order are zero (e.g., additive models and pairwise-interaction models), but also include other models, such as nearest-neighbor interaction models. Projecting onto the hierarchical gauge (but not other parametric family gauges) is guaranteed to produce a parameter vector where the appropriate entries are still fixed to be zero.

#### Zero-sum gauge

The zero-sum gauge (illustrated in Figs. 1A and 2A) is the hierarchical gauge for which *p* is the uniform distribution. The name of this gauge comes from the fact that, when *p* is uniform, Eq. (24) becomes

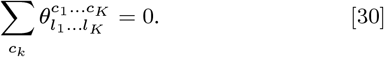

Prior studies (12, 15) have characterized the zero-sum gauge for the all-order interaction model. Our formulation of the hierarchical gauge extends those findings and generalizes them to gauges defined by non-uniformly weighted sums of parameters.

#### Wild-type and generalized wild-type gauges

The wild-type gauge (illustrated in Figs. 1B and 2B) is a hierarchical gauge that arises in the limit as *p* approaches an indicator function for some “wild-type sequence,” *s*^*wt*^. In the wild-type gauge, only the parameters θ_*s*_^*′*^ for which *s*^*′*^ matches *s*^*wt*^ receive any penalization, and all these penalized θ_*s*_^*′*^(except for θ_0_) are driven to zero. Consequently, θ_0_ quantifies the activity of the wild-type sequence, each 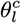 quantifies the effect of a single mutation to the wild-type sequence, each 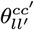 quantifies the epistatic effect of two mutations to the wild-type sequence, and so on. However, seeing the wild-type gauge as a special case of the hierarchical gauge provides the possibility of generalizing the wild-type gauge by using a *p* that is not the indicator function on a single sequence but rather defines a distribution over one or more alleles per position that can be considered as being “wild-type” (equivalently, the frequencies of some subset of position-specific characters are set to zero). These gauges all inherit the property from the the hierarchical gauge that their coefficients relate to the average effect of taking draws from the probability distribution defined by *p* and setting a subset of positions to the characters specified by that coefficient. More rigorously, these gauges are defined by considering the limit lim_ϵ*→*0_^+^ of the hierarchical gauge with factorizable distribution

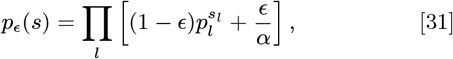

where the 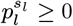 are the position-specific factors of the desired nonnegative vector of probabilities *p*.

### Applications

We now demonstrate the utility of our results on two example models of complex sequence-function relationships. First, we study how the parameters of the all-order interaction model behave under different parametric gauges in the context of a simulated landscape on short binary sequences. We observe that model parameters exhibit nontrivial collective behavior across different choices of gauge. Second, we examine the parameters of an empirical pairwise-interaction model for protein GB1 using the zero-sum and multiple generalized wild-type gauges. We observe how these different hierarchical gauges enable different interpretations of model parameters and facilitate the derivation of simplified models that are approximately correct in different localized regions of sequence space. The results provide intuition for the behavior of the various parametric gauges, and show in particular how hierarchical gauges can be used to explore and interpret real sequence-function relationships.

#### Gauge-fixing a simulated landscape on short binary sequences

To illustrate the consequences of choosing gauges in the parametric family, we consider a simulated random landscape on short binary sequences. Consider sequences of length *L* = 3 built from the alphabet 𝒜 = *{*0, 1}, and assume that the activities of these sequences are as shown in Fig. 3A. The corresponding all-order interaction model has (α + 1)^*L*^ = 27 parameters, which we index using augmented sequences: 1 constant parameter (θ_*\*\*\**_), 6 additive parameters (θ_0*\*\**_, θ_1*\*\**_, θ_*\**0*\**_, θ_*\**1*\**_, θ_*\*\**0_, θ_*\*\**1_), 12 pairwise parameters (θ_00*\**_, θ_01*\**_, θ_10*\**_, θ_11*\**_, θ_0*\**0_, θ_0*\**1_, θ_1*\**0_, θ_1*\**1_, θ_*\**00_, θ_*\**01_, θ_*\**10_, θ_*\**11_), and 8 third-order parameters (θ_000_, θ_001_, θ_010_, θ_011_, θ_100_, θ_101_, θ_110_, θ_111_).

**Fig. 3.**
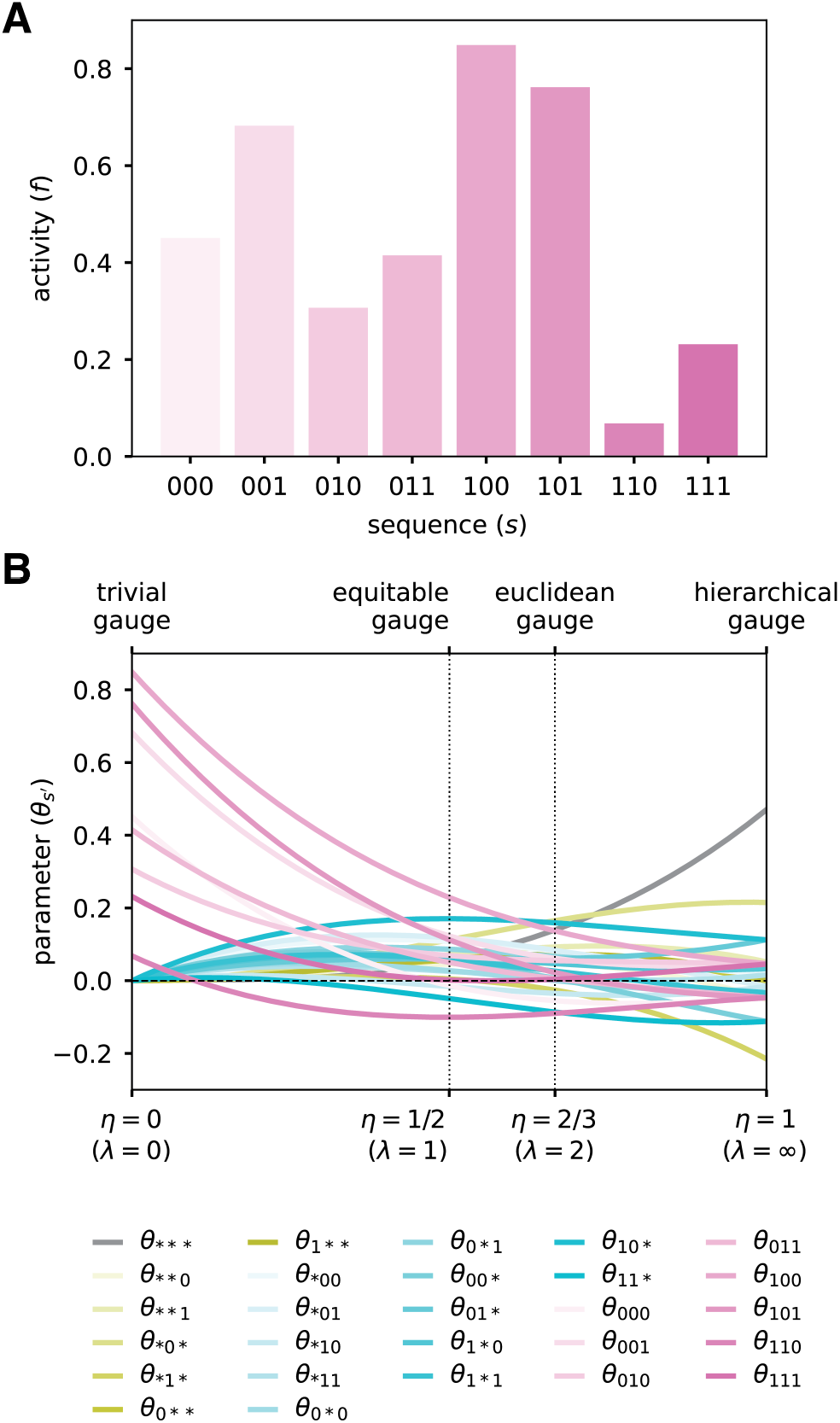
Binary landscape expressed in various parametric family gauges. (A) Simulated random activity landscape for binary sequences of length *L* = 3. (B) Parameters of the all-order interaction model for the binary landscape as functions of *η* = *λ/*(1 + *λ*). Values of *η* corresponding to different named gauges are indicated. Note: because the uniform distribution is assumed in all these gauges, the hierarchical gauge is also the zero-sum gauge.

We now consider what happens to the values of these 27 parameters when they are expressed in different parametric gauges, Θ^*λ*,*p*^. Specifically, we assume that *p* is the uniform distribution and vary the parameter *λ* from 0 to ∞ (equivalent, η varies from 0 to 1). Note that each entry in the projection matrix *P*^*λ*,*p*^ (Eq. 19) is a cubic function of η, due to *L* = 3. Consequently, each of the 27 gauge-fixed model parameters is a cubic function of η [Fig. 3B]. In the trivial gauge (λ = 0, η = 0), only the 8 third-order parameters are nonzero, and the values of the 8 third-order parameters correspond to the values of the landscape at the 8 corresponding sequences. In the equitable gauge (λ = 1, η = 1/2), the spread of the 8 third-order parameters about zero is larger than that of the 12 pairwise parameters, which is larger than that of the 6 additive parameters, which is larger than that of the constant parameter. In the euclidean gauge (λ = 2, η = 2/3), the parameters of all orders exhibit a similar spread about zero. In the hierarchical gauge (λ = ∞, η = 1), the spread of the 8 third-order parameters about zero is smaller than that of the 12 pairwise parameters, which is smaller than that of the 6 additive parameters, which is smaller than that of the constant parameter. Moreover, the marginalization and orthogonality properties of the hierarchical gauge fix certain parameters to be equal or opposite to each other, e.g. we must have θ_1*\*\**_ = −θ_0*\*\**_ and the third order parameters are all equal up to their sign, which depends only on whether the corresponding sequence feature has an even or odd number of “1”s.

This example illustrates generic features of the parametric gauges. For any all-order interaction model on sequences of length *L*, the entries of the projection matrix *P*^*λ*,*p*^ will be *L*-order polynomials in η. Consequently, the values of model parameters, when expressed in the gauge Θ^*λ*,*p*^, will also be *L*-order polynomials in η. In the trivial gauge, only the highest-order parameters will be nonzero. In the equitable gauge, the spread about zero will tend to be smaller for lower-order parameters relative to higher-order parameters. In the euclidean gauge, parameters of all orders will exhibit similar spread about zero. In the zero-sum gauge, the spread about zero will tend to be minimized for higher-order parameters relative to lower-order parameters. The nontrivial quantitative behavior of model parameters in different parametric gauges thus underscores the importance of choosing a specific gauge before quantitatively interpreting parameter values.

#### Hierarchical gauges of an empirical landscape for protein GB1

Projecting model parameters onto different hierarchical gauges can facilitate the exploration and interpretation of sequence-function relationships. To demonstrate this application of gauge fixing, we consider an empirical sequence-function relationship describing the binding of the GB1 protein to immunoglobulin G (IgG). Wu et al. (59) performed a deep mutational scanning experiment that measured how nearly all 20^4^ = 160, 000 amino acid combinations at positions 39, 40, 41, and 54 of GB1 affect GB1 binding to IgG. These data report log_2_ enrichment values for each assayed sequence relative to the wild-type sequence at these positions, VDGV (Fig. 4A,B). Using these data and least-squares regression, we inferred a pairwise interaction model for log_2_ enrichment as a function of protein sequence at these *L* = 4 variable positions. The resulting pairwise interaction model comprises 1 constant parameter, 80 additive parameters, and 2400 pairwise parameters. Fig. S1 illustrates the performance of this model. To understand the structure of the activity landscape described by the pairwise interaction model, we now examine the values of model parameters in multiple hierarchical gauges. Explicit formulas for implementing hierarchical gauges for pairwise-interaction models are given in SI Sec. 8.

**Fig. 4.**
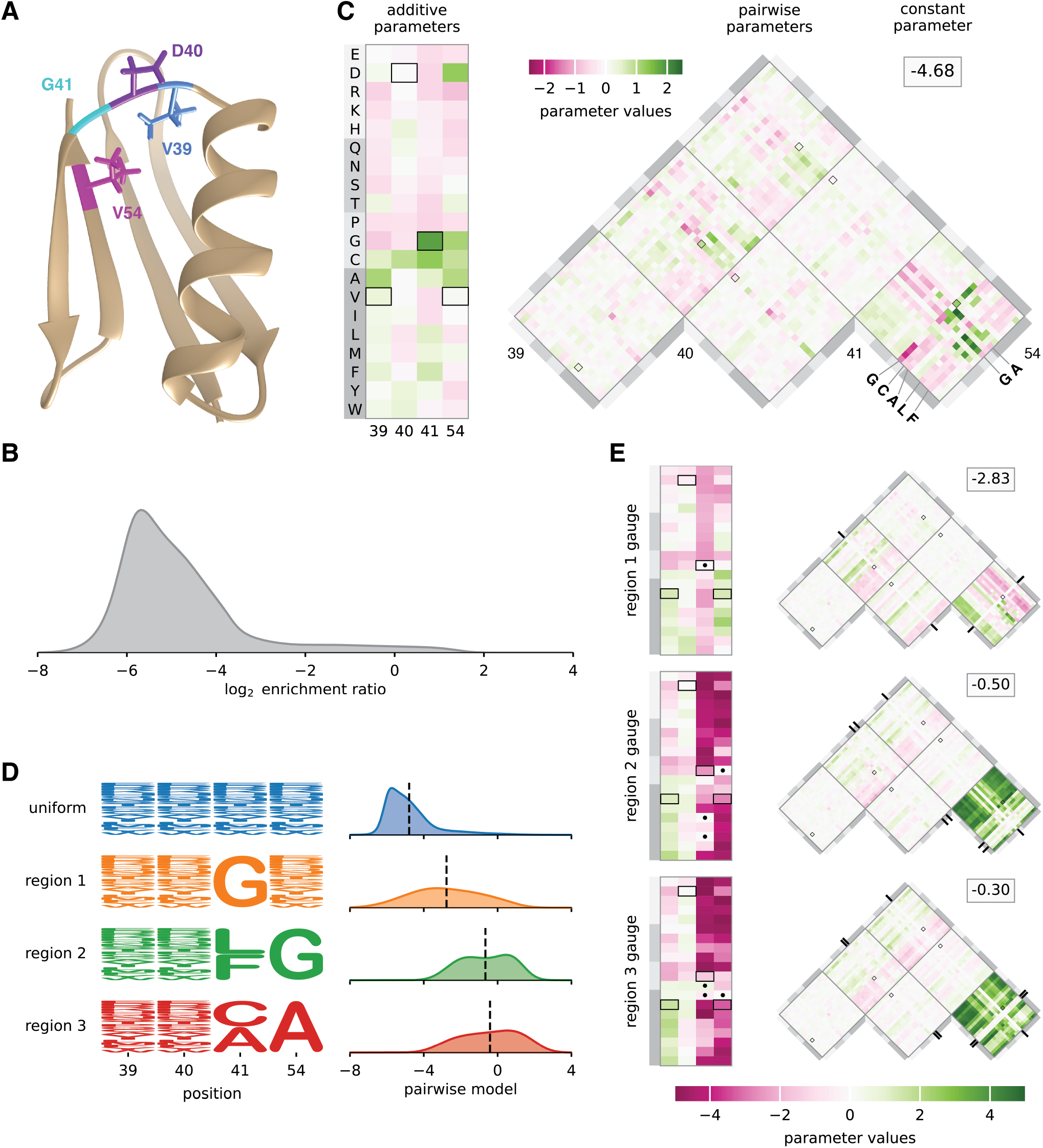
Landscape exploration using hierarchical gauges. (A) NMR structure of GB1, with residues V39, D40, G41, and V54 shown [PDB: 3GB1, from (58)]. (B) Distribution of log_2_ enrichment values measured by (59) for nearly all 160,000 GB1 variants having mutations at positions 39, 40, 41, and 54. (C) Pairwise interaction model parameters inferred from the data of (59), expressed in the uniform hierarchical gauge (i.e., the zero-sum gauge). Boxes indicate parameters contributing to the wild-type sequence, VDGV. (D) Probability logos (57) for uniform, region 1, region 2, and region 3 sequence distributions. Distributions of pairwise interaction model predictions for each region are also shown. (E) Model parameters expressed in the region 1, region 2, and region 3 hierarchical gauges. Dots and tick marks indicate region-specific constraints. Probability densities (panels B and D) were estimated using DEFT (41). Pairwise interaction model parameters were inferred by least-squares regression using MAVE-NN (57). Regions 1, 2, and 3 were defined based on (60). NMR: nuclear magnetic resonance. GB1: domain B1 of protein G.

Fig. 4C shows the parameters of the pairwise interaction model expressed in the hierarchical gauge corresponding to a uniform probability distribution on sequence space (i.e., the zero-sum gauge). In the zero-sum gauge, the constant parameter θ_0_ equals the average activity of all sequences. We observe θ_0_ = −4.68, indicating that a typical random sequence is depleted approximately 20-fold relative to the wild-type sequence, which the pairwise interaction model assigns a score of −.21. This finding confirms the expectation that a random sequence should be substantially less functional than the wildtype sequence.

The additive parameters in the zero-sum gauge are shown in the rectangular heat map in Fig. 4C, and each additive parameter is equal to the difference between the mean activity of the set of sequences containing the corresponding amino acid at the relevant position relative to the mean activity of random sequences. We observe that the wild-type sequence receives positive or near-zero contributions at every position, including a contribution from the most positive additive parameter, corresponding to G at position 41. The additive parameters at positions 39, 40, and 54 that contribute to the wild-type sequence, however, are not the largest additive parameters at these positions. Moreover, the additive parameters that contribute to the wild-type sequence only sum to 2.32, meaning that, of the total difference (4.47) between the wild-type sequence score and the average sequence score, almost half (2.15) is due to contributions from pairwise parameters. This finding quantifies the importance of epistatic interactions at positions 39, 40, 41, and 54 for the IgG binding activity of wild-type GB1.

The pairwise parameters in the zero-sum gauge are shown in the triangular heat map in Fig. 4C, where each pairwise parameter is equal to the difference between the observed mean of the sequences containing the specified pair of characters at the specified pair of conditions and the expected mean activity based on the the mean activity of sequences containing the individual characters and the grand mean activity. We observe that the three largest-magnitude pairwise contributions to the wildtype sequence are from the pair G41V54 (1.25), V39G41 (0.91), and D40G41 (-0.44), indicating that position 41 is a major hub of epistatic interactions contributing to the wild-type sequence. Moving to the landscape as a whole, we observe that the largest magnitude pairwise interactions link positions 41 and 54. Moreover, the strongest positive pairwise contributions are obtained when a small amino acid (G or A) is present at position 54, and a G, C, A, L, or P is present at position 41 (see also 45). This finding provides insight into the chemical nature of the epistatic interactions that facilitate wild-type GB1 binding to IgG.

Previous work (60, 61) identified three disjoint regions of high-activity sequences (region 1, region 2, and region 3) in the GB1 landscape measured by Wu et al. (59). Region 1 comprises sequences with G at 41; region 2 comprises sequences with L or F at position 41 and G at position 54; and region 3 comprises sequences with C or A at position 41 and A at position 54. To investigate the structure of the GB1 landscape within the three regions, we defined probability distributions that were uniform in each region of sequence space and zero outside (Fig. 4D; see SI Sec. 8 for formal definitions of these regions). We then examined the values of the parameters of the pairwise-interaction model, with the parameters expressed in the hierarchical gauges corresponding to the probability distribution *p*(*s*) for each of the three regions (the “region 1 hierarchical gauge”, “region 2 hierarchical gauge”, and “region 3 hierarchical gauge”). Since some characters at positions 41 and 54 have had their frequencies set to zero, these hierarchical gauges are in fact generalized wild-type gauges, and the additive and pairwise parameters can be interpreted in terms of the mean effects of introducing mutations to these specific regions of sequences space.

In the region 1 hierarchical gauge (Fig. 4E, top), the additive parameters for position 41 quantify the effect of mutations away from G, and the additive parameters for positions 39, 40, and 54 quantify the average effect of mutations conditional on G at position 41. From the additive parameters at position 54, we observe that cysteine (C) and hydrophobic residues (A, V, I, L, M, or F) increase binding, and that proline (P) and charged residues (E, D, R, K) decrease binding. From the additive parameters at position 40, we observe that amino acids with a 5-carbon or 6-carbon ring (H, F, Y, W) increase binding, suggesting the presence of structural constraints on side chain shape, rather than constraints on hydrophobicity or charge. The largest pairwise parameters all involve mutations from G at position 41 to another amino acid, and careful inspection of these pairwise parameters show that the pairwise parameters are roughly equal and opposite to the additive effects of mutations at the other three positions. This indicates a classical form of masking epistasis, where the typical effect of a mutation at position 41 results in a more or less complete loss of function, after which mutations at the remaining three positions no longer have a substantial effect .

In the region 2 hierarchical gauge (Fig. 4E, middle), the additive parameters at position 54 quantify the average effect of mutations away from G contingent on L or F at position 41, the additive parameters at position 41 quantify the average effects of mutations away from L or F contingent on G at position 54, and the additive parameters at positions 39 and 40 quantify the average effects of mutations contingent on L or F at position 41 and on G at position 54. From the values of the additive parameters, we observe that mutations away from L or F at position 41 in the presence of G at position 54 are typically strongly deleterious (mean effect -3.39), and that mutations away from G at position 54 in the presence of L or F at position 41 are also strongly deleterious (mean effect -3.75). However, the pairwise parameters linking positions 41 and 54 are strongly positive (mean effect 2.85), again indicating a masking effect where the first deleterious mutation at position 41 or 54 results in a more or less complete loss of function, so that an additional mutation at the other position has little effect (note the similar but less extreme pattern of masking between the large effect mutations at positions 41 and 54 with the milder mutations at positions 40 and 41, whose interaction coefficients are of the opposite sign of the additive effects at positions 40 and 41). Similar results hold for the region 3 hierarchical gauge, where mutations at positions 41 and 54 have masking effects on each other as well as on mutations in the other two positions (Fig. 4E, bottom). However, we can also contrast patterns of mutational effects between these regions. For example, mutating position 54 to G (a mututation leading towards region 2) on average has little effect in region 1 but would be deleterious in region 3. Similarly, if we consider mutations leading from region 2 to region 3, we can see that mutating 41 to C in region 2 typically has little effect whereas mutating 41 to A is more deleterious .

Besides using the interpretation of hierarchical gauge parameters as average effects of mutations to understand how mutational effects differ in different regions of sequence space, we hypothesised that by applying different hierarchical gauges to the pairwise interaction model, one might be able to obtain simple additive models that are accurate in different regions of sequence space. Our hypothesis was motivated by the fact that the parameters of all-order interaction models in the zero-sum gauge are chosen to maximize the fraction of variance in the sequence-function relationship that is explained by lower-order parameters. To test our hypothesis, we defined an additive model for each of the four hierarchical gauges described above (uniform, region 1, region 2, and region 3) by projecting pairwise interaction model parameters onto the hierarchical gauge for that region then setting all the pairwise parameters to zero. We then evaluated the predictions of each additive model on sequences randomly drawn from each of the four corresponding probability distributions (uniform, region 1, region 2, and region 3). The results (Fig. 5) show that the activities of sequences sampled uniformly from the sequence space are best explained by the additive model derived from the zero-sum gauge, that the activities of region 1 sequences are best explained by the additive model derived from the region 1 hierarchical gauge, and so on for regions 2 and 3. This shows that projecting a pairwise interaction model (or other hierarchical one-hot model) onto the hierarchical gauge corresponding to a specific region of sequence space can sometimes be used to obtain simplified models that approximate predictions by the original model in that region.

**Fig. 5.**
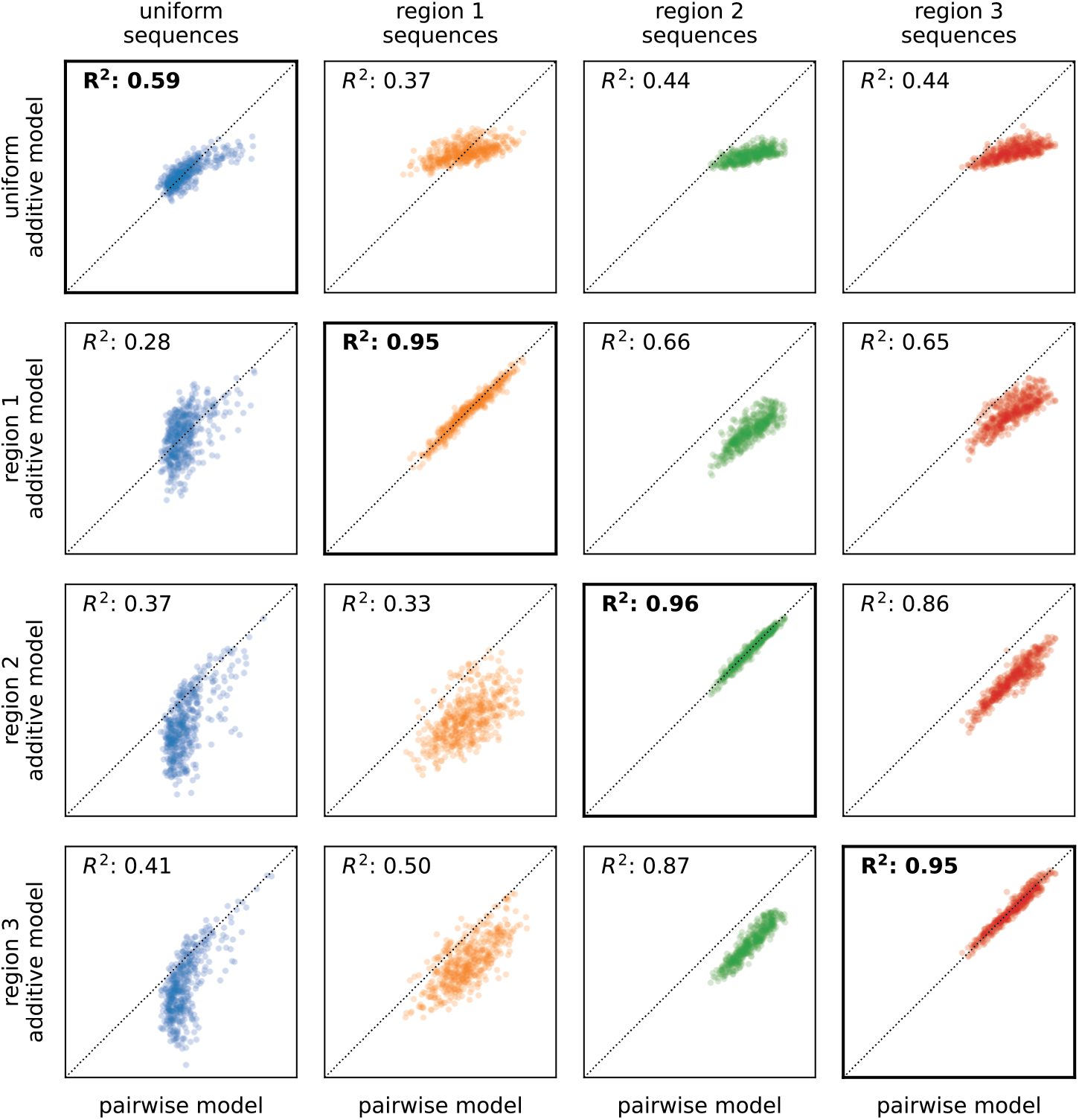
Model coarse-graining using hierarchical gauges. Predictions of additive models for GB1 derived by model truncation using region-specific zero-sum gauges (from Fig. 4C,E), plotted against predictions of the full pairwise-interaction model, are shown for 500 sequences randomly sampled from each of the four distributions listed in Fig. 4D (i.e., uniform, region 1, region 2, and region 3). Diagonals indicate equality. GB1: domain B1 of protein G.

## Discussion

Here we report a unified strategy for fixing the gauge of commonly used models of sequence-function relationships. First we defined a family of analytically tractable gauges for the all-order interaction model. We then derived explicit formulae for imposing any of these gauges on model parameters, and used these formulae to investigate the mathematical properties of the these gauges. The results show that these gauges include multiple commonly used gauges, and that a subset of these gauges (the hierarchical gauges) can be applied to diverse lower-order models (including additive models, pairwise-interaction models, and higher-order interaction models).

Next, we demonstrated the family of gauges in two contexts: a simulated all-order interaction landscape on short binary sequences, and an empirical pairwise-interaction landscape for the protein GB1. The GB1 results, in particular, show how applying different hierarchical gauges can facilitate the biological interpretation of complex models of sequence-function relationships and to derive simplified models that are approximately correct in localized regions of sequence space.

Our study was limited to linear models of sequence-function relationships. Although linear models are used in many computational biology applications, more complex models are becoming increasingly common. For example, linear-nonlinear models [which include global epistasis models (9, 62–64) and thermodynamic models (56, 57, 65–68)] are commonly used to describe fitness landscapes and/or sequence-dependent biochemical activities. In addition to the gauge freedoms of their linear components, linear-nonlinear models can have additional gauge freedoms, such as diffeomorphic modes (69, 70), that also need to be fixed before parameter values can be meaningfully interpreted.

Sloppy modes are another important issue to address when interpreting quantitative models of sequence-function relationships. Sloppy modes are directions in parameter space that (unlike gauge freedoms) do affect model predictions but are nevertheless poorly constrained by data (71, 72). Understanding the mathematical structure of sloppy modes, and developing systematic methods for fixing these modes, is likely to be more challenging than understanding gauge freedoms. This is because sloppy modes arise from a confluence of multiple factors: the mathematical structure of a model, the distribution of data in feature space, and measurement uncertainty. Nevertheless, understanding sloppy modes is likely to be as important in many applications as understanding gauge freedoms. We believe the study of sloppy modes in quantitative models of sequence-function relationships is an important direction for future research.

Deep neural network (DNN) models present perhaps the biggest challenge for parameter interpretation. DNN models have had remarkable success in quantitatively modeling biological sequence-function relationships, most notably in the context of protein structure prediction (73, 74), but also in the context of other processes including gene regulation (75–77), epigenetics (78–80), and mRNA splicing (81, 82). It remains unclear, however, how researchers might gain insights into the molecular mechanisms of biological processes from inferred DNN models. DNNs are by nature highly over-parameterized (83–85), making the direct interpretation of DNN parameters infeasible. Instead, a variety of attribution methods have been developed to facilitate DNN model interpretations (86– 89). Existing attribution methods can often be thought of as providing additive models that approximate DNN models in localized regions of sequence space (90), and the presence of gauge freedoms in these additive models needs to be addressed when interpreting attribution method output [as in (91, 92)]. We anticipate that, as DNN models become more widely adopted for mechanistic studies in biology, there will be a growing need for attribution methods that provide more complex quantitative models that approximate DNN models in localized regions of sequence space (16). If so, a comprehensive mathematical understanding of gauge freedoms in parametric models of sequence-function relationships will be needed to aid in these DNN model interpretations.

## Materials and Methods

See Supplemental Information detailed derivations of mathematical results. All data and Python scripts used to generate the figures are available at https://github.com/jbkinney/23_posfai.

## ACKNOWLEDGMENTS

We thank Peter Koo for helpful conversations and Samantha Petti for comments on the manuscript. This work was supported by NIH grant R35 GM133613 (AP, JZ, DMM), NIH grant R35 GM133777 (AP, JBK), NIH grant R01 HG011787 (JBK), the Alfred P. Sloan foundation (DMM), as well as additional funding from the Simons Center for Quantitative Biology at CSHL (DMM, JBK) and the College of Liberal Arts and Sciences at the University of Florida (JZ).

## Notes

https://github.com/jbkinney/23_posfai

### Competing Interest Statement

The authors have declared no competing interest.

### Summary of Updates

We have added an acknowledgement of helpful comments from Samantha Petti.

