## Supplemental Information for "Gauge fixing for sequence-function relationships"

### Supplemental Information for:
### Gauge fixing for sequence-function relationships

Anna Posfai, Juannan Zhou, David M. McCandlish, Justin B. Kinney

June 22, 2024

#### Contents


1 Sequences and embeddings 1

2 Gauge freedoms 1

3 Linear gauges 2

4 All-order interaction models 5

5 Parametric family of gauges 8

6 Trivial gauge, euclidean gauge, and equitable gauge 13

7 Hierarchical gauges 14

8 Hierarchical gauges of an empirical landscape for protein GB1 19

9 Appendix 20


#### 1 Sequences and embeddings

**Definition 1.** *The alphabet $\mathcal{A}$ is an ordered set of $\alpha$ characters $(c_1, \ldots, c_\alpha)$.*

**Definition 2.** *Sequence space $\mathcal{S}$ is the set of all sequences of length $L$ built from characters in the alphabet $\mathcal{A}$. We use $N$ to denote the number of sequences in $\mathcal{S}$, and $s_l$ to denote the character at position $l$ in sequence $s$.*

**Definition 3.** *An embedding $\vec{x}$ is a mapping from $\mathcal{S}$ to a real vector space $V = \mathbb{R}^M$. We use $M$ throughout to denote the dimension of $V$.*

**Definition 4.** *The embedding space $S$ of an embedding $\vec{x} : \mathcal{S} \to V$ is a vector space defined by $S \equiv \mathrm{span}(\{\vec{x}(s) : s \in \mathcal{S}\})$. $S$ is also called the span of $\vec{x}$.*

**Definition 5.** *The design matrix $X$ of an embedding $\vec{x} : \mathcal{S} \to V$ is an $N \times M$ matrix having elements $X_{ij} = [\vec{x}(s_i)]_j$, where $i = 1, \ldots, N$ indexes all sequences in $\mathcal{S}$ (the specific order does not matter) and $j = 1, \ldots, M$ indexes the dimensions of $V$.*

#### 2 Gauge freedoms

**Definition 6.** *The space of gauge freedoms (a.k.a. freedom space) $G$ of an embedding $\vec{x} : \mathcal{S} \to V$ is a vector space defined by*

$$G \equiv \left\{ \vec{g} \in V : \vec{g}^{\top} \vec{x}(s) = 0 \ \forall \ s \in \mathcal{S} \right\}. \tag{1}$$

*We use $\gamma$ to denote the dimension of $G$.*

**Claim 1.** *Let $\vec{x}$ be an embedding, $S$ be the embedding space of $\vec{x}$, and $G$ be the freedom space of $\vec{x}$. Then $G$ is the orthogonal complement of $S$, i.e., $G = S^{\perp}$.*

*Proof.* First we show that $G \subseteq S^\perp$. Consider any $\vec{g} \in G$. By Definition 6, $\vec{g}^\top \vec{x}(s) = 0$ for every $s \in \mathcal{S}$, and so $\vec{g}$ is orthogonal to $S$, and thus $\vec{g} \in S^\perp$. This establishes that $G \subseteq S^\perp$. Next we show that $S^\perp \subseteq G$. If $\vec{v} \in S^\perp$, then $\vec{v}^\top \vec{w} = 0$ for every $\vec{w} \in S$. In particular, $\vec{v}^\top \vec{x}(s) = 0$ for every $s \in \mathcal{S}$, implying that $\vec{v} \in G$. This establishes that $S^\perp \subseteq G$, proving the claim. □

**Gauge freedoms of pairwise-interaction models.** Eq. 7 in the main text appears to gives $2\alpha$ linear relations for every pair of positions $l < l'$, namely

$$x_l^c(s) = \sum_{c' \in \mathcal{A}} x_{ll'}^{cc'}(s) \quad (\alpha \text{ linear relations}), \tag{2}$$

$$x_{l'}^{c'}(s) = \sum_{c \in \mathcal{A}} x_{ll'}^{cc'}(s) \quad (\alpha \text{ linear relations}). \tag{3}$$

However, these linear relations are not independent, as summing Eq. 2 over all $c \in \mathcal{A}$ gives the same linear relation as summing Eq. 3 over all $c' \in \mathcal{A}$:

$$1 = \sum_{c,c' \in \mathcal{A}} x_{ll'}^{cc'}(s). \tag{4}$$

Eq. 4 is in fact the only dependency between the $2\alpha$ linear relations in Eq. 2 and Eq. 3. Therefore, there are actually $2\alpha - 1$ independent gauge freedoms per pair of positions $l < l'$, and there are $\binom{L}{2}$ such pairs of positions. We thus find a total of

$$\gamma_{\text{pairwise}} = L + \binom{L}{2}(2\alpha - 1) \tag{5}$$

gauge freedoms for the pairwise-interaction model.

### 3 Linear gauges

**Definition 7.** *A linear gauge space $\Theta$ for an embedding $\vec{x} : \mathcal{S} \to V$ having freedom space $G$ is a vector space such that, for all $\vec{v} \in V$, there is a unique decomposition $\vec{v} = \vec{\theta} + \vec{g}$ where $\vec{\theta} \in \Theta$ and $\vec{g} \in G$. All gauge spaces discussed in what follows are assumed to be linear. Note that $\dim \Theta = M - \gamma$.*

**Definition 8.** *A projection matrix $P$ that projects from a vector space $V$ into a vector space $V_1$ along a vector space $V_2$ is a matrix such that, for all $\vec{v} \in V$, $P\vec{v} \in V_1$ and $\vec{v} - P\vec{v} \in V_2$.*

**Claim 2.** *Let $\vec{x} : \mathcal{S} \to V$ be an embedding, let $G$ be the freedom space of $\vec{x}$, and let $\Theta$ be a gauge space of $\vec{x}$. Then there is a unique a projection matrix $P$ that projects $V$ into $\Theta$ along $G$. Moreover, the matrix $Q \equiv I_M - P$ is the unique matrix that projects $V$ into $G$ along $\Theta$.*

*Proof.* Let $\vec{e}_1, \ldots, \vec{e}_\gamma$ be a basis for $G$, and $\vec{f}_1, \ldots, \vec{f}_{M-\gamma}$ be a basis for $\Theta$. Using these basis vectors as columns, define the $M \times \gamma$ matrix $E \equiv (\vec{e}_1, \ldots, \vec{e}_\gamma)$ and the $M \times (M - \gamma)$ matrix $F \equiv (\vec{f}_1, \ldots, \vec{f}_{M-\gamma})$. Choose any $\vec{v} \in V$. By Definition 7, $\vec{v}$ can be uniquely decomposed as

$$\vec{v} = \vec{\theta} + \vec{g} \tag{6}$$

where $\vec{\theta} \in \Theta$ and $\vec{g} \in G$. Because the columns of $E$ and $F$ provide bases for $G$ and $\Theta$, respectively, there is a $\gamma$-dimensional vector $\vec{a}$ and an $(M - \gamma)$-dimensional vector $\vec{b}$ such that $\vec{g} = E\vec{a}$ and $\vec{\theta} = F\vec{b}$. Therefore,

$$\vec{v} = E\vec{a} + F\vec{b} \tag{7}$$

$$= (E \;\; F) \begin{pmatrix} \vec{a} \\ \vec{b} \end{pmatrix} \tag{8}$$

$$\Rightarrow \begin{pmatrix} \vec{a} \\ \vec{b} \end{pmatrix} = (E \;\; F)^{-1} \vec{v} \tag{9}$$

$$\Rightarrow \vec{\theta} = (0_{M\times\gamma} \;\; F) \begin{pmatrix} \vec{a} \\ \vec{b} \end{pmatrix} \tag{10}$$

$$= (0_{M\times\gamma} \;\; F)\,(E \;\; F)^{-1} \vec{v}, \tag{11}$$

where $(E \;\; F)$ is the $M \times M$ matrix given by horizontally concatenating $E$ and $F$, and $0_{M\times\gamma}$ is an $M \times \gamma$ matrix of zeros. A projection matrix $P$ from $V$ into $\Theta$ along $G$ therefore exists and is given by

$$P = (0_{M\times\gamma} \;\; F)\,(E \;\; F)^{-1}. \tag{12}$$

To see that $Q$ projects $V$ into $G$ along $\Theta$, simply note that, for any $\vec{v} \in V$, $Q\vec{v} = \vec{v} - P\vec{v} \in G$ and $\vec{v} - Q\vec{v} = P\vec{v} \in \Theta$. To prove that $P$ is unique, assume that there is another matrix $P' \neq P$ that projects into $\Theta$ along $G$. There must therefore be a $\vec{v} \in V$ such that $P'\vec{v} \neq P\vec{v}$. By Definition 8, $P'\vec{v} \in \Theta$ and $\vec{v} - P'\vec{v} \in G$. But $P\vec{v} \in \Theta$ and $\vec{v} - P\vec{v} \in G$ as well, and by Definition 7, the decomposition of $\vec{v}$ into a component in $\Theta$ and a component in $G$ is unique, implying that $P\vec{v} = P'\vec{v}$, which is a contradiction. An analogous proof shows that $Q$ is unique. □

**Definition 9.** *A metric $\Lambda$ on an $M$-dimensional vector space $V$ is a symmetric positive-definite $M \times M$ matrix. The $\Lambda$-inner product of two vectors $\vec{v}, \vec{w} \in V$ is defined to be*

$$\langle \vec{v}, \vec{w} \rangle_\Lambda \equiv \vec{v}^\top \Lambda \vec{w}. \tag{13}$$

*The $\Lambda$-norm of a vector $\vec{v} \in V$ is defined to be*

$$||\vec{v}||_\Lambda \equiv \sqrt{\langle \vec{v}, \vec{v} \rangle_\Lambda} = \sqrt{\vec{v}^\top \Lambda \vec{w}}. \tag{14}$$

*$\vec{v}$ and $\vec{w}$ are $\Lambda$-orthogonal if and only if $\langle \vec{v}, \vec{w} \rangle_\Lambda = 0$.*

**Definition 10.** *An orthogonalizing metric, $\Lambda$, for a gauge space $\Theta$ and freedom space $G$ is a metric for which all $\vec{g} \in G$ are $\Lambda$-orthogonal to all $\vec{\theta} \in \Theta$.*

**Claim 3.** *Let $\vec{x} : \mathcal{S} \to V$ be an embedding, $G$ be the freedom space of $\vec{x}$, and $\Theta$ be a gauge space of $\vec{x}$. Then there exists an orthogonalizing metric $\Lambda$ for $\Theta$ and $G$.*

*Proof.* Let $P$ be the projection matrix from $V$ into $\Theta$ along $G$, let $Q = I_M - P$, and define the matrix $\Lambda \equiv P^\top P + Q^\top Q$. It is a simple matter to show that $\Lambda$ is symmetric and positive-definite, and is thus a metric. Using $P^2 = P$, $Q^2 = Q$ and $Q = I_M - P$, it is readily shown that $PQ$ are both equal to the zero matrix and hence $\Lambda$ satisfies $\Lambda = P^\top \Lambda P + Q^\top \Lambda Q$. Therefore, by Claim 25 in the Appendix, $\Lambda$ orthogonalizes $\Theta$ and $G$. □

**Claim 4.** *Let $\vec{x} : \mathcal{S} \to V$ be an embedding, $X$ be the design matrix of $\vec{x}$, $G$ be the freedom space of $\vec{x}$, $\Theta$ be a gauge space of $\vec{x}$, and $\Lambda$ be a metric that orthogonalizes $\Theta$ and $G$. Then for any vector $\vec{\theta}_{\text{init}} \in V$, the vector $\vec{\theta}^*$ in the gauge orbit of $\vec{\theta}_{\text{init}}$ that has minimal $\Lambda$-norm lies in $\Theta$.*

*Proof.* Let $\theta^*$ be the unique element of $\Theta$ that lies in the gauge orbit of $\vec{\theta}_{\text{init}}$. Because $\Lambda$ orthogonalizes $\Theta$ and $G$, $\vec{g}^\top \Lambda \vec{\theta}^* = 0$ for all $\vec{g} \in G$. By Claim 25,

$$||\vec{\theta}^* + \vec{g}||^2_\Lambda \geq ||\vec{\theta}^*||^2_\Lambda, \tag{15}$$

with equality obtaining only when $\vec{g} = \vec{0}$. This proves that $\vec{\theta}^*$ is the unique vector with the smallest $\Lambda$-norm of any vector in the gauge orbit of $\vec{\theta}_{\text{init}}$. □

**Claim 5.** *Let $\vec{x} : \mathcal{S} \to V$ be an embedding, $X$ be the design matrix of $\vec{x}$, $G$ be the freedom space of $\vec{x}$, $\Theta$ be a gauge space of $\vec{x}$, and $\Lambda$ be a metric that orthogonalizes $\Theta$ and $G$. Then the matrix $P$ that projects along $G$ into $\Theta$ is given by $P = \Lambda^{-1/2}(X\Lambda^{-1/2})^+ X$.*

*Proof.* Consider the transformed embedding $\vec{x}' = \Lambda^{-1/2}\vec{x}$, which corresponds to the transformed design matrix $X' = X\Lambda^{-1/2}$ as well as the transformed embedding space $S' = \Lambda^{-1/2}S$. If a parameter vector $\vec{\theta} \in V$ yields model predictions $X\vec{\theta}$), then $\vec{\theta}' = \Lambda^{1/2}\vec{\theta}$ yields the same model predictions $X'\vec{\theta}'$. Consequently, $G' = \Lambda^{1/2}G$ is the transformed freedom space and $\Theta' = \Lambda^{-1/2}\Theta$ is the transformed gauge space. Because $\Theta$ and $G$ are $\Lambda$-orthogonal, $\Theta'$ and $G'$ are orthogonal in the Euclidean sense. The transformed embedding space $S'$ is also orthogonal to $G'$, and so $\Theta' = S'$ The matrix $P'$ that projects along $G'$ and onto $\Theta'$ is therefore the orthogonal projection matrix onto the space spanned by the rows of $X'$, and is given by $P' = (X')^+ X$. Now consider an initial parameter $\vec{\theta}_{\text{init}} \in V$, its gauge-fixed counterpart $\vec{\theta}_{\text{fixed}} = P\vec{\theta}_{\text{init}} \in \Theta$ as well as the transformed versions of these vectors, $\vec{\theta}'_{\text{init}} = \Lambda^{-1/2}\vec{\theta}_{\text{init}}$ and $\vec{\theta}'_{\text{fixed}} = \Lambda^{-1/2}\vec{\theta}_{\text{fixed}} \in \Theta'$. One can readily verify that

$$X\vec{\theta}_{\text{init}} = X\vec{\theta}_{\text{fixed}} = X'\vec{\theta}'_{\text{init}} = X'\vec{\theta}'_{\text{fixed}}. \tag{16}$$

Therefore,

$$\begin{aligned}
\vec{\theta}_{\text{fixed}} &= \Lambda^{-1/2}\vec{\theta}'_{\text{fixed}}, && (17)\\
&= \Lambda^{-1/2}P'\vec{\theta}'_{\text{init}}, && (18)\\
&= \Lambda^{-1/2}(X')^+ X'\vec{\theta}'_{\text{init}}, && (19)\\
&= \Lambda^{-1/2}(X\Lambda^{-1/2})^+ X\Lambda^{-1/2}\Lambda^{1/2}\vec{\theta}_{\text{init}}, && (20)\\
&= P\vec{\theta}_{\text{init}} && (21)
\end{aligned}$$

where $P = \Lambda^{-1/2}(X\Lambda^{-1/2})^+ X$. This proves the claim. □

**Definition 11.** *A loss function $\mathcal{L}$ is said to be $L_2$-regularized by a metric $\Lambda$ iff it has the form*

$$\mathcal{L}(\vec{\theta}) = \mathcal{L}_{\text{data}}(X\vec{\theta}) + \frac{\beta}{2}\vec{\theta}^\top \Lambda \vec{\theta}, \tag{22}$$

*where $X$ is the design matrix of an embedding $\vec{x}$, $\mathcal{L}_{\text{data}}$ is a data-dependent loss function that depends only on model predictions $X\vec{\theta}$, and $\beta$ is a positive scalar.*

**Claim 6.** *Let $\vec{x} : \mathcal{S} \to V$ be an embedding, $G$ be the freedom space of $\vec{x}$, $\Theta$ be a gauge space of $\vec{x}$, $\Lambda$ be a metric that orthogonalizes $\Theta$ and $G$, $\mathcal{L}$ be a loss function that is $L_2$-regularized by $\Lambda$, and $\vec{\theta}^*$ be a minimum of $\mathcal{L}$. Then $\vec{\theta}^* \in \Theta$.*

*Proof.*

$$\begin{aligned}
\vec{\theta}^* &= \operatorname{argmin}_{\vec{v}\in V}\ \mathcal{L}(\vec{v}) && (23)\\
&= \operatorname{argmin}_{\vec{v}\in V}\ \left[\mathcal{L}_{\text{data}}(X\vec{v}) + \frac{\beta}{2}\vec{v}^\top\Lambda\vec{v}\right] && (24)\\
&= \operatorname{argmin}_{\left\{\vec{v}=\vec{\theta}+\vec{g}:\vec{\theta}\in\Theta,\,\vec{g}\in G\right\}}\ \left[\mathcal{L}_{\text{data}}(X(\vec{\theta}+\vec{g})) + \frac{\beta}{2}(\vec{\theta}+\vec{g})^\top\Lambda(\vec{\theta}+\vec{g})\right] && (25)\\
&= \operatorname{argmin}_{\vec{\theta}\in\Theta}\ \left[\mathcal{L}_{\text{data}}(X\vec{\theta}) + \frac{\beta}{2}\vec{\theta}^\top\Lambda\vec{\theta}\right] + \operatorname{argmin}_{\vec{g}\in G}\ \left[\frac{\beta}{2}\vec{g}^\top\Lambda\vec{g}\right] && (26)\\
&= \operatorname{argmin}_{\vec{\theta}\in\Theta}\ \left[\mathcal{L}_{\text{data}}(X\vec{\theta}) + \frac{\beta}{2}\vec{\theta}^\top\Lambda\vec{\theta}\right]. && (27)
\end{aligned}$$

In Eq. 25 we used the fact that $\vec{v} \in V$ can be expressed uniquely as $\vec{v} = \vec{\theta} + \vec{g}$ for $\vec{\theta} \in \Theta$ and $\vec{g} \in G$ (by Definition 7), and in Eq. 26 we used the fact that $X\vec{g} = \vec{0}$ for any $\vec{g} \in G$ (by Definition 6), together with the assumption that $\Lambda$ orthogonalizes $\Theta$ and $G$. We thus find that $\vec{\theta}^* \in \Theta$. □

**Claim 7.** *Let $\vec{x} : \mathcal{S} \to V$ be an embedding, $G$ be the freedom space of $\vec{x}$, $\Theta$ be a gauge space of $\vec{x}$, $\mathcal{L}_{\text{data}}$ be a data-dependent loss function that depends only on model predictions $X\vec{\theta}$, and $\Delta$ be an $N \times N$ positive definite matrix. Then there exists a metric $\Lambda$ defining a loss $\mathcal{L}$ that is $L_2$-regularized by $\Lambda$ that has the following property: every minimum $\vec{\theta}^*$ of $\mathcal{L}$ lies within $\Theta$ and satisfies*

$$\mathcal{L}(\theta^*) = \mathcal{L}_{\text{data}}(X\vec{\theta}^*) + \frac{\beta}{2}\left(X\vec{\theta}^*\right)^\top \Delta \left(X\vec{\theta}^*\right). \tag{28}$$

*Proof.* By Claim 6 it suffices to construct a $\Lambda$ that is an orthogonalizing metric for $\Theta$ and $G$ and which satisfies Equation 28. Let $P$ be the projection matrix from $V$ into $\Theta$ along $G$, let $Q = I_M - P$, and define the matrix $\Lambda \equiv P^\top X^\top \Delta X P + Q^\top Q$. It is a simple matter to show that $\Lambda$ is symmetric and positive-definite, and is thus a metric. Using $P^2 = P$ and $Q^2 = Q$, and $Q = I_M - P$, it is readily shown that $PQ$ are both equal to the zero matrix and hence $\Lambda$ satisfies $\Lambda = P^\top \Lambda P + Q^\top \Lambda Q$. Therefore, by Claim 25 in the Appendix, $\Lambda$ orthogonalizes $\Theta$ and $G$. Then by Claim 6, we have $\vec{\theta}^* \in \Theta$ and hence $P\vec{\theta}^* = \vec{\theta}^*$ and $\vec{\theta}^*$ is in the null space of $Q$. Consequently,

$$\begin{aligned}
\mathcal{L}(\theta^*) &= \mathcal{L}_{\text{data}}(X\vec{\theta}^*) + \frac{\beta}{2}\vec{\theta}^{*\top}\left(P^\top X^\top \Delta X P + Q^\top Q\right)\vec{\theta}^* && (29)\\
&= \mathcal{L}_{\text{data}}(X\vec{\theta}^*) + \frac{\beta}{2}\left(\vec{\theta}^{*\top}P^\top X^\top \Delta X P\vec{\theta}^* + \vec{\theta}^{*\top}Q^\top Q\vec{\theta}^*\right) && (30)\\
&= \mathcal{L}_{\text{data}}(X\vec{\theta}^*) + \frac{\beta}{2}\left(X\vec{\theta}^*\right)^\top \Delta \left(X\vec{\theta}^*\right) && (31)
\end{aligned}$$

as required. □

**Practical recipe for fixing a linear gauge using $L_2$ regularization.** Claim 7 shows that for any choice of linear gauge $\Theta$ and any desired positive definite $L_2$ regularizer $\Delta$ on model predictions, we can construct a positive definite $L_2$ regularizer $\Lambda$ on model parameters that penalizes the vector of predictions according to $\Delta$ and results in an inferred parameter vector $\vec{\theta}^*$ that is guaranteed to be a member of our desired gauge $\Theta$. Practically, the steps to calculate $\Lambda$ are:

1. Find a basis $\vec{e}_1, \ldots, \vec{e}_\gamma$ for the null space of the design matrix $X$. This can be done, for instance, by using Gaussian elimination to column reduce the block matrix $\begin{pmatrix} X \\ I_M \end{pmatrix}$, where the resulting matrix will have $\gamma$ non-zero columns whose top $N$ entries are 0 and whose bottom $N$ entries will each serve as one vector in the desired basis. See see ref. [1] for analytical methods to determine this basis for the all-order interaction model and related models.

2. Find a basis $\vec{f}_1, \ldots, \vec{f}_{M-\gamma}$ for the desired linear gauge space $\Theta$. Any linearly independent set of $M - \gamma$ vectors that are members of $\Theta$ will suffice.

3. Using these basis vectors as columns, define the $M \times \gamma$ matrix $E \equiv (\vec{e}_1, \ldots, \vec{e}_\gamma)$ and the $M \times (M-\gamma)$ matrix $F \equiv (\vec{f}_1, \ldots, \vec{f}_{M-\gamma})$. Then calculate the projection matricies $P = (0_{M\times\gamma}\ F)\,(E\ \ F)^{-1}$ and $Q = I_M - P$.
4. Set $\Lambda \equiv P^\top X^\top \Delta X P + Q^\top Q$.
5. Minimize $\mathcal{L}(\vec{\theta}) \equiv \mathcal{L}_{\text{data}}(X\vec{\theta}) + \frac{\beta}{2}\vec{\theta}^\top \Lambda \vec{\theta}$ for some choice of regularization parameter $\beta > 0$.

Note similarly that given a specified $L_2$ regularizer $\Lambda$ on model parameters, the induced $L_2$ regularizer on model predictions is given by $\Delta \equiv (PX^+)^\top \Lambda\, (PX^+)$ which satisfies $x^\top \Delta x > 0$ for all nonzero $x$ in the column space of $X$.

### 4 All-order interaction models

**Definition 12.** *The position-specific augmented embedding $\vec{x}'_l : \mathcal{S} \to V_l$, where $V_l = \mathbb{R}^{\alpha+1}$, is given by*

$$\vec{x}'_l(s) = \begin{pmatrix} x^*_l(s) \\ x^{c_1}_l(s) \\ \cdots \\ x^{c_\alpha}_l(s) \end{pmatrix} \quad \text{for all} \quad s \in \mathcal{S}, \tag{32}$$

*where, as in the main text, $x^*_l(s) = 1$ for all $s \in \mathcal{S}$ and, for all $c \in \mathcal{A}$, $x^c_l(s)$ is one if $s_l = c$ and is zero otherwise. In what follows, we use $G_l$ to denote the freedom space of $\vec{x}'_l$ and $S_l$ to denote the embedding space of $\vec{x}'_l$.*

**Claim 8.** *$S_l$ has dimension $\alpha$ and is given by $\left\{(\beta^*, \beta^{c_1}, \ldots, \beta^{c_\alpha})^\top : \beta^* = \sum_{c\in\mathcal{A}} \beta^c\right\}$; $G_l$ has dimension $1$ and is spanned by the vector $(-1, 1, \ldots, 1)^\top$.*

*Proof.* Let $S_l$ denote the span of $\vec{x}'_l$. By definition, any vector $\vec{v} \in S_l$ can be written as a linear combination of vectors $\vec{x}'_l(s)$ over $s \in S$, i.e.,

$$\vec{v} = \sum_{s\in\mathcal{S}} \beta(s) \begin{pmatrix} x^*_l(s) \\ x^{c_1}_l(s) \\ \vdots \\ x^{c_\alpha}_l(s) \end{pmatrix} \tag{33}$$

for some mapping $\beta : \mathcal{S} \to \mathbb{R}$. Defining $\beta^c \equiv \sum_{s\in\mathcal{S}} \beta(s) x^c_l(s)$ for all $c \in \mathcal{A}'$, we get

$$\vec{v} = \begin{pmatrix} \beta^* \\ \beta^{c_1} \\ \vdots \\ \beta^{c_\alpha} \end{pmatrix}. \tag{34}$$

The $\alpha + 1$ values $\{\beta^c\}_{c\in\mathcal{A}}$ are arbitrary except for one constraint arising from the fact that $x^*_l(s) = \sum_{c\in\mathcal{A}} x^c_l(s)$ for all $s \in \mathcal{S}$:

$$\begin{aligned} \sum_{c\in\mathcal{A}} \beta^c &= \sum_{c\in\mathcal{A}} \sum_{s\in\mathcal{S}} \beta(s) x^c_l(s) && (35) \\ &= \sum_{s\in\mathcal{S}} \beta(s) \sum_{c\in\mathcal{A}} x^c_l(s) && (36) \\ &= \sum_{s\in\mathcal{S}} \beta(s) x^*_l(s) && (37) \\ &= \beta^*. && (38) \end{aligned}$$

We therefore see that $S_l$ has dimension $\alpha$ and comprises the vectors stated in Claim 8.

Now let $\vec{g} = (-1, 1, \ldots, 1)^\top$. Taking the dot product of $\vec{g}$ with $\vec{x}'_l$ gives

$$\vec{g}^\top \vec{x}'(s) = -x^*_l(s) + \sum_{c\in\mathcal{A}} x^c_l(s) = 0 \quad \text{for all} \quad s \in \mathcal{S}, \tag{39}$$

This shows that $\vec{g} \in G_l$ and, in light of the fact that $\dim G_l = \dim V - \dim S_l = 1$, proves that $G_l$ is spanned by $\vec{g}$. □

**Definition 13.** *The all-order embedding $\vec{x}_{\text{all}} : \mathcal{S} \to V_{\text{all}}$ ($V_{\text{all}} = \mathbb{R}^M$ where $M = (\alpha+1)^L$) is defined by the tensor product*

$$\vec{x}_{\text{all}}(s) = \bigotimes_{l=1}^{L} \vec{x}'_l(s) \quad \text{for all} \quad s \in \mathcal{S}. \tag{40}$$

*We use $S_{\text{all}}$ to denote the embedding space of $\vec{x}_{\text{all}}$, and $G_{\text{all}}$ to denote the freedom space of $\vec{x}_{\text{all}}$.*

**Claim 9.** *The embedding space of $\vec{x}_{\text{all}}$ is given by*

$$S_{\text{all}} = \bigotimes_{l=1}^{L} S_l. \tag{41}$$

*Proof.* For each $c \in \mathcal{A}$, define $\alpha$ distinct $(\alpha+1)$-dimensional vectors $\vec{e}_c$, one for each $c \in \mathcal{A}$ and with elements indexed by $c' \in \mathcal{A}'$ given by

$$[\vec{e}_c]_{c'} = \begin{cases} 1 & \text{if } c' = * \text{ or } c' = c \\ 0 & \text{otherwise} \end{cases} \tag{42}$$

Note that, for any $s \in \mathcal{S}$, $\vec{e}_{s_l} = \vec{x}'_l(s)$. Next, for each position $l$, define the set of vectors $\mathcal{B}_l = \{\vec{e}_c\}_{c\in\mathcal{A}}$. From the proof of Claim 8, $\mathcal{B}_l$ forms a linearly independent basis for $S_l$. By definition of the tensor product of vector spaces, a basis for the space $S \equiv \bigotimes_{l=1}^{L} S_l$ is given by

$$\begin{aligned}
\mathcal{B} &\equiv \left\{ \bigotimes_{l=1}^{L} \vec{v}_l : \vec{v}_1 \in \mathcal{B}_1, \ldots, \vec{v}_L \in \mathcal{B}_L \right\} && (43)\\
&= \left\{ \bigotimes_{l=1}^{L} \vec{e}_{c_l} : c_1, \ldots, c_L \in \mathcal{A} \right\} && (44)\\
&= \left\{ \bigotimes_{l=1}^{L} \vec{e}_{s_l} : s \in \mathcal{S} \right\} && (45)\\
&= \left\{ \bigotimes_{l=1}^{L} \vec{x}'_l(s) : s \in \mathcal{S} \right\} && (46)\\
&= \{\vec{x}_{\text{all}}(s) : s \in \mathcal{S}\}. && (47)
\end{aligned}$$

$\mathcal{B}$ is therefore also a basis for $S_{\text{all}}$. Since $S$ and $S_{\text{all}}$ share the same basis, they are the same vector space. Note that we have learned, in the process, that all vectors $\vec{x}_{\text{all}}(s)$, $s \in \mathcal{S}$, are linearly independent. □

**Claim 10.** *The freedom space of $\vec{x}_{\text{all}}$ is given by*

$$G_{\text{all}} = \bigoplus_{(R_1,\ldots,R_L)\in\mathcal{R}} \left[ \bigotimes_{l=1}^{L} R_l \right], \tag{48}$$

*where*

$$\mathcal{R} = \{(R_1, \ldots, R_L) : R_l \in \{S_l, G_l\} \textit{ for all } l \textit{ and } R_l = G_l \textit{ for at least one } l\}. \tag{49}$$

*Proof.*

$$\begin{aligned}
V_{\text{all}} &= \bigotimes_{l=1}^{L} \mathrm{V}_l && (50)\\
&= \bigotimes_{l=1}^{L} [S_l \oplus G_l] && (51)\\
&= \left[ \bigotimes_{l=1}^{L} S_l \right] \oplus \left[ \bigoplus_{\{R_1,\ldots,R_L\}\in\mathcal{R}} \bigotimes_{l=1}^{L} R_l \right] && (52)\\
&= S_{\text{all}} \oplus W && (53)
\end{aligned}$$

where

$$W \equiv \bigoplus_{\{R_1,\ldots,R_L\}\in\mathcal{R}} \bigotimes_{l=1}^{L} R_l, \tag{54}$$

This shows that $\dim W = \dim V_{\text{all}} - \dim S_{\text{all}} = \dim G_{\text{all}}$. However, this does not yet show that $W = G_{\text{all}}$.

To see that $W = G_{\text{all}}$, let $\{\vec{u}_l^c\}_{c\in\mathcal{A}'}$ be a basis for $V_l$. Each $\vec{u}_l^c$ has a unique decomposition $\vec{u}_l^c = \vec{\theta}_l^c + \vec{g}_l^c$ where $\vec{\theta}_l^c \in S_l$ and $\vec{g}_l^c \in G_l$. For every augmented sequence $s' \in \mathcal{S}'$, there is a corresponding vector in $V_{\text{all}}$ given by $\vec{u}_{s'} = \bigotimes_{l=1}^{L} \vec{u}_l^{s_l'}$. Moreover, the set $\{\vec{u}_{s'}\}_{s'\in\mathcal{S}'}$ is a basis for $V_{\text{all}}$. The vector $\vec{u}_{s'}$ can be decomposed into a component in $S_{\text{all}}$ and a component in $W$ via

$$\begin{aligned}
\vec{u}_{s'} &= \bigotimes_{l=1}^{L} \vec{u}_l^{s'_l} &(55)\\
&= \bigotimes_{l=1}^{L} \left[\vec{\theta}_l^{s'_l} + \vec{g}_l^{s'_l}\right] &(56)\\
&= \bigotimes_{l=1}^{L} \vec{\theta}_l^{s'_l} + \sum_{(\vec{r}_1,\ldots,\vec{r}_L)\in\rho_{s'}} \bigotimes_{l=1}^{L} \vec{r}_l &(57)\\
&= \vec{\theta}_{s'} + \vec{r}_{s'} &(58)
\end{aligned}$$

where

$$\begin{aligned}
\vec{\theta}_{s'} &\equiv \bigotimes_{l=1}^{L} \vec{\theta}_l^{s'_l}, &(59)\\
\vec{r}_{s'} &\equiv \sum_{(\vec{r}_1,\ldots,\vec{r}_L)\in\rho_{s'}} \bigotimes_{l=1}^{L} \vec{r}_l, &(60)\\
\rho_{s'} &\equiv \left\{(\vec{r}_1,\ldots,\vec{r}_L) : \vec{r}_l \in \left\{\vec{\theta}_l^{s'_l}, \vec{g}_l^{s'_l}\right\} \text{ for all } l \text{, and } \vec{r}_l = \vec{g}_l^{s'_l} \text{ for at least one } l\right\}. &(61)
\end{aligned}$$

Observe that $r_{s'} \in W$ for every $s' \in \mathcal{S}'$. Moreover, since $\{\vec{u}_{s'}\}_{s'\in\mathcal{S}'}$ is a basis for $\vec{V}_{\text{all}}$, $\{\vec{r}_{s'}\}_{s'\in\mathcal{S}'}$ is a basis for $W$. Finally, observe that for every $s \in \mathcal{S}$, the dot product of $\vec{x}_{\text{all}}(s)$ with $\vec{r}_{s'}$ is zero, i.e.,

$$[\vec{x}_{\text{all}}(s)]^\top \vec{r}_{s'} = \sum_{(\vec{r}_1,\ldots,\vec{r}_L)\in\rho_{s'}} \prod_{l=1}^{L} [\vec{x}'_l(s)]^\top \vec{r}_l = 0, \tag{62}$$

because $\vec{r}_l \in G_l$ for at least one $l$ in each product. Therefore, $\vec{r}_{s'} \in G_{\text{all}}$ for every $s' \in \mathcal{S}'$. Since $\{\vec{r}_{s'}\}_{s'\in\mathcal{S}'}$ is a basis for $W$, we conclude that $W = G_{\text{all}}$.

□

**Claim 11.** *For each position $l$, let $\Theta_l$ be a gauge space for $\vec{x}'_l$, let $P_l$ be projection matrix into $\Theta_l$ along $G_l$, and let $\Lambda_l$ be a metric that orthogonalizes $\Theta_l$ and $G_l$. Then $\Theta \equiv \bigotimes_l \Theta_l$ is a gauge space of $\vec{x}_{\text{all}}$, $P \equiv \bigotimes_l P_l$ projects into $\Theta$ along $G_{\text{all}}$, and $\Lambda \equiv \bigotimes_l \Lambda_l$ is a metric that orthogonalizes $\Theta$ and $G_{\text{all}}$.*

*Proof.* For each position $l = 1,\ldots,L$, let $\vec{v}_l = \vec{\theta}_l + \vec{g}_l$ where $\vec{v}_l \in V_l$, $\vec{\theta}_l \in \Theta_l$, and $\vec{g}_l \in G_l$. Defining $\vec{v} = \bigotimes_{l=1}^{L} \vec{v}_l$, we get

$$\vec{v} = \bigotimes_{l=1}^{L} \left[\vec{\theta}_l + \vec{g}_l\right] = \vec{\theta} + \vec{g} \tag{63}$$

where $\vec{\theta} = \bigotimes_{l=1}^{L} \vec{\theta}_l$ lives in the space $\Theta \equiv \bigotimes_{l=1}^{L} \Theta_l$, and

$$\begin{aligned}
\vec{g} &= \sum_{(\vec{r}_1,\ldots,\vec{r}_L)\in\rho} \bigotimes_{l=1}^{L} \vec{r}_l, &(64)\\
\rho &\equiv \left\{(\vec{r}_1,\ldots,\vec{r}_L) : \vec{r}_l \in \left\{\vec{\theta}_l, \vec{g}_l\right\} \text{ for all } l \text{, and } \vec{r}_l = \vec{g}_l \text{ for at least one } l\right\}, &(65)
\end{aligned}$$

lives in the space

$$G \equiv \bigoplus_{(R_1,\ldots,R_L)\in\mathcal{R}} \left[\bigotimes_{l=1}^{L} R_l\right], \tag{66}$$

where

$$\mathcal{R} = \{(R_1,\ldots,R_L) : R_l \in \{\Theta_l, G_l\} \text{ for all } l \text{ and } R_l = G_l \text{ for at least one } l\}. \tag{67}$$

By arguments similar to those in Claim 10, it is readily seen that $\vec{x}_{\text{all}}(s)^\top \vec{g} = \vec{0}$ for every $s \in \mathcal{S}$. Moreover, by arguments similar to those in Claim 10, it is readily seen that this construction is able to yield a basis of vectors $\vec{g}$ for $G$, proving that that $G = G_{\text{all}}$ is the freedom space of $\vec{x}_{\text{all}}$. Therefore, for every vector of the form $\vec{v} = \bigotimes_{l=1}^{L} \vec{v}_l$, there is a unique decomposition $\vec{v} = \vec{\theta} + \vec{g}$, where $\vec{\theta} \in \Theta$ and $\vec{g} \in G_{\text{all}}$. Because a basis of such vectors $\vec{v}$ can be found for $V$, we see that any vector $\vec{v} \in V$ can be uniquely decomposed as $\vec{v} = \vec{\theta} + \vec{g}$, where $\vec{\theta} \in \Theta$ and $\vec{g} = G_{\text{all}}$. This proves that $\Theta$ is a gauge space of $\vec{x}_{\text{all}}$.

Defining $P \equiv \bigotimes_{l=1}^{L} P_l$ and applying this to $\vec{v}$, we find that

$$P\vec{v} = \bigotimes_{l=1}^{L} P_l \vec{v}_l = \bigotimes_{l=1}^{L} \vec{\theta}_l = \vec{\theta} \tag{68}$$

is in $\Theta$, which implies that $\vec{v} - P\vec{v} = \vec{g}$ is in $G_{\text{all}}$. Since a basis of vectors $\vec{v}$ for $V$ can be found that decompose in this way, all vectors in $V$ decompose in this manner. This proves that $P$ projects into $\Theta$ along $G_{\text{all}}$

Now define $\Lambda \equiv \bigotimes_{l=1}^{L} \Lambda_l$. It is readily seen that $\Lambda$ is symmetric and positive definite from the fact that every $\Lambda_l$ is symmetric and positive definite. Moreover,

$$\begin{aligned}
\Lambda &= \bigotimes_{l=1}^{L} (P_l^\top \Lambda_l P_l + Q_l^\top \Lambda_l Q_l) && (69)\\
&= \bigotimes_{l=1}^{L} P_l^\top \Lambda_l P_l + \sum_{(R_1,\ldots,R_L)\in\mathcal{R}} \bigotimes_{l=1}^{L} R_l^\top \Lambda_l R_l && (70)\\
&= \left[\bigotimes_{l=1}^{L} P_l\right]^\top \left[\bigotimes_{l=1}^{L} \Lambda_l\right] \left[\bigotimes_{l=1}^{L} P_l\right] + \sum_{(R_1,\ldots,R_L)\in\mathcal{R}} \left[\bigotimes_{l=1}^{L} R_l\right]^\top \left[\bigotimes_{l=1}^{L} \Lambda_l\right] \left[\bigotimes_{l=1}^{L} R_l\right] && (71)\\
&= \left[\bigotimes_{l=1}^{L} P_l\right]^\top \left[\bigotimes_{l=1}^{L} \Lambda_l\right] \left[\bigotimes_{l=1}^{L} P_l\right] + \left[\sum_{(R_1,\ldots,R_L)\in\mathcal{R}} \bigotimes_{l=1}^{L} R_l\right]^\top \left[\bigotimes_{l=1}^{L} \Lambda_l\right] \left[\sum_{(R_1,\ldots,R_L)\in\mathcal{R}} \bigotimes_{l=1}^{L} R_l\right] && (72)\\
&= P^\top \Lambda P + Q^\top \Lambda Q. && (73)
\end{aligned}$$

where $Q = I_M - P$ and the set $\mathcal{R}$ of ordered operator sets is defined to be

$$\mathcal{R} \equiv \{(R_1,\ldots,R_L) : R_i = P_i \text{ or } R_i = Q_i \text{ for all } i = 1,\ldots,L, \text{ with } R_i = Q_i \text{ for at least one } i\}. \tag{74}$$

In Eq. 69, we used the fact that $\Lambda_l$ satisfies

$$\Lambda_l = P_l^\top \Lambda_l P_l^\top + Q_l^\top \Lambda_l Q_l, \tag{75}$$

where $Q_l \equiv I_{(\alpha+1)} - P_l$ (see Claim 25). In going from Eq. 71 to Eq. 72, we used the fact that, if two ordered sets of operators $(R_1,\ldots,R_L)$ and $(R'_1,\ldots,R'_L)$ in $\mathcal{R}$ are different, then

$$\left[\bigotimes_{l=1}^{L} R_l\right]^\top \left[\bigotimes_{l=1}^{L} \Lambda_l\right] \left[\bigotimes_{l=1}^{L} R'_l\right] = \bigotimes_{l=1}^{L} R_l^\top \Lambda R'_l = 0 \tag{76}$$

since there will be an $l$ such that $R_l^\top \Lambda_l R'_l$ is either $P_l^\top \Lambda_l Q_l = 0$ or $Q_l \Lambda_l P_l = 0$. In going from Eq. 72 to Eq. 73, we used the fact that

$$\sum_{(R_1,\ldots,R_L)\in\mathcal{R}} \bigotimes_{l=1}^{L} R_l = \bigotimes_{i=1}^{L} (P_l + Q_l) - \bigotimes_{i=1}^{L} P_l = I_M - P = Q. \tag{77}$$

The fact that $P$ projects into $\Theta$ along $G_{\text{all}}$, together with the result $\Lambda = P^\top \Lambda P + Q^\top \Lambda Q$ in Eq. 73 proves (by Claim 25) that $\Lambda$ orthogonalizes $\Theta$ and $G_{\text{all}}$.

□

### 5 Parametric family of gauges

**Definition 14.** *A probability distribution $p$ on $\mathcal{S}$ is positive iff $p(s) > 0$ for all $s \in \mathcal{S}$.*

**Definition 15.** *A factorizable probability distribution $p$ on $\mathcal{S}$ is one that can be written $p(s) = \prod_{l=1}^{L} p_l^{s_l}$ for all $s \in \mathcal{S}$, where $p_l$ is a probability distribution over the $\alpha$ possible characters at position $l$, i.e., $p(s) = p_l^{s_l}$.*

**Definition 16.** *An augmented sequence $s'$ is a sequence built from the augmented alphabet $\mathcal{A}' = \{*, c_1, \ldots, c_\alpha\}$, where $c_1, \ldots, c_\alpha$ are the characters in $\mathcal{A}$ and $*$ is a wild-card character that is interpreted as matching any character in $\mathcal{A}$. The set of all augmented sequences is denoted $\mathcal{S}'$. Note that every augmented sequence $s'$ is can be interpreted as a subset of $\mathcal{S}$ that comprises sequences matching the pattern defined by $s'$. For this reason we will sometimes use expressions like $s \in s'$ and $s' \subseteq t'$ (for $s \in \mathcal{S}$ and $s', t' \in \mathcal{S}$).*

To aid in our discussion of the all-order interaction model [Eq. **??**], we define an augmented alphabet $\mathcal{A}' = \{*, c_1, \ldots, c_\alpha\}$, where $c_1, \ldots, c_\alpha$ are the characters in $\mathcal{A}$ and $*$ is a wild-card character that is interpreted as matching any character in $\mathcal{A}$. Let $\mathcal{S}'$ denote the set of sequences of length $L$ comprising characters from $\mathcal{A}'$. For each augmented sequence $s' \in \mathcal{S}'$, we define the sequence feature $x_{s'}(s)$ to be 1 if a sequence $s$ matches the pattern described by $s'$ and to be 0 otherwise. In this way, each augmented sequence $s'$ serves as a regular expression against which bona fide sequences are compared.

**Definition 17.** *Given a non-negative real number $\lambda$, a factorizable probability distribution $p$ on $\mathcal{S}$, and a sequence position $l$, the position-specific parametric gauge $\Theta_l^{\lambda,p}$ is defined as*

$$\Theta_l^{\lambda,p} \equiv V_\lambda \oplus V_\perp^{p_l}, \tag{78}$$

*where $V_\lambda \equiv \text{span } \{(\lambda, 1, \cdots, 1)^\top\}$ and $V_\perp^{p_l} \equiv \left\{(0, v_{c_1}, \ldots, v_{c_\alpha})^\top : \sum_{c \in \mathcal{A}} p_l^c v_c = 0\right\}$.*

**Claim 12.** *The matrix $P_l^{\lambda,p}$ that projects $V_l$ along $G_l$ and onto $\Theta_l^{\lambda,p}$ is an $(\alpha+1) \times (\alpha+1)$ matrix given by*

$$P_l^{\lambda,p} = \begin{pmatrix} \eta & p_l^{c_1}\eta & \cdots & p_l^{c_\alpha}\eta \\ 1-\eta & 1-p_l^{c_1}\eta & \cdots & -p_l^{c_\alpha}\eta \\ \vdots & \vdots & \ddots & \vdots \\ 1-\eta & -p_l^{c_1}\eta & \cdots & 1-p_l^{c_\alpha}\eta \end{pmatrix}, \tag{79}$$

*where $\eta \equiv \lambda/(1+\lambda)$.*

*Proof.* Any $(\alpha+1)$-dimensional vector $\vec{v} \in V_l$ can be decomposed as

$$\vec{v} = \vec{\theta} + \vec{g}, \tag{80}$$

where $\vec{\theta} \in \Theta_l^{\lambda,p}$ and $\vec{g} \in G_l$. From the definitions of $\Theta_l^{\lambda,p}$ and $G$, $\vec{\theta}$ and $\vec{g}$ must have the forms

$$\vec{\theta} = c_\lambda \vec{v}_\lambda + \vec{\theta}_\perp \quad \text{and} \quad \vec{g} = c_{-1}\vec{e}_{-1}, \tag{81}$$

where

$$\vec{e}_\lambda = \begin{pmatrix} \lambda \\ 1 \\ \vdots \\ 1 \end{pmatrix}, \quad \vec{e}_{-1} = \begin{pmatrix} -1 \\ 1 \\ \vdots \\ 1 \end{pmatrix}, \quad \vec{\theta}_\perp = \begin{pmatrix} 0 \\ \theta_l^{c_1} \\ \vdots \\ \theta_l^{c_\alpha} \end{pmatrix}, \tag{82}$$

and where $c_\lambda$ and $c_{-1}$ are real numbers that depend on $\vec{v}$. The projection matrix $Q_l^{\lambda,p} \equiv I - P_l^{\lambda,p}$ projects $V_l$ into $G_l$ along $\Theta_l^{\lambda,p}$, and so is related to the scalar $c_{-1}$ via

$$Q_l^{\lambda,p}\vec{v} = c_{-1}\vec{e}_{-1}. \tag{83}$$

We now compute $c_{-1}$ as a function of $\vec{v}$. First we define the metric

$$\Lambda \equiv \begin{pmatrix} 1 & 0 & \cdots & 0 \\ 0 & \alpha p_l^{c_1} & \cdots & 0 \\ \vdots & \vdots & \ddots & \vdots \\ 0 & 0 & \cdots & \alpha p_l^{c_\alpha} \end{pmatrix}, \tag{84}$$

and compute

$$\vec{e}_\lambda^\top \Lambda \vec{e}_\lambda = \lambda^2 + \sum_{i=1}^{\alpha} \alpha p_l^{c_i} = \alpha + \lambda^2, \tag{85}$$

$$\vec{e}_{-1}^\top \Lambda \vec{e}_\lambda = -\lambda + \sum_{i=1}^{\alpha} \alpha p_l^{c_i} = \alpha - \lambda, \tag{86}$$

$$\vec{e}_{-1}^{\top}\Lambda\vec{e}_{-1} = 1+\sum_{i=1}^{\alpha}\alpha p_l^{c_i} = \alpha+1, \tag{87}$$

$$\vec{e}_{\lambda}^{\top}\Lambda\vec{\theta}_{\perp} = 0+\sum_{i=1}^{\alpha}\alpha p_l^{c_i}\theta_l^{c_i} = 0, \tag{88}$$

$$\vec{e}_{-1}^{\top}\Lambda\vec{\theta}_{\perp} = 0+\sum_{c}\sum_{i=1}^{\alpha}\alpha p_l^{c_i}\theta_l^{c_i} = 0. \tag{89}$$

Next we solve for $c_{-1}$ via

$$\begin{aligned}
\vec{e}_{\lambda}^{\top}\Lambda\vec{v} &= c_\lambda(\vec{e}_{\lambda}^{\top}\Lambda\vec{e}_\lambda)+c_{-1}(\vec{e}_{\lambda}^{\top}\Lambda\vec{e}_{-1})+(\vec{e}_{\lambda}^{\top}\Lambda\vec{\theta}_{\perp}) && (90)\\
&= c_\lambda(\alpha+\lambda^2)+c_{-1}(\alpha-\lambda). && (91)\\
\vec{e}_{-1}^{\top}\Lambda\vec{v} &= c_\lambda(\vec{e}_{-1}^{\top}\Lambda\vec{e}_\lambda)+c_{-1}(\vec{e}_{-1}^{\top}\Lambda\vec{e}_{-1})+(\vec{e}_{-1}^{\top}\Lambda\vec{\theta}_{\perp}) && (92)\\
&= c_\lambda(\alpha-\lambda)+c_{-1}(\alpha+1). && (93)\\
\Rightarrow (\alpha-\lambda)\vec{e}_{\lambda}^{\top}\Lambda\vec{v}-(\alpha+\lambda^2)\vec{e}_{-1}^{\top}\Lambda\vec{v} &= c_{-1}\left[(\alpha-\lambda)^2-(\alpha+1)(\alpha+\lambda^2)\right] && (94)\\
&= c_{-1}\left[-2\alpha\lambda-\alpha(1+\lambda^2)\right] && (95)\\
&= -\alpha(1+2\lambda+\lambda^2)c_{-1} && (96)\\
&= -\alpha(1+\lambda)^1 c_{-1} && (97)\\
\Rightarrow c_{-1} &= -\frac{(\alpha-\lambda)}{\alpha(1+\lambda)^2}\vec{e}_{\lambda}^{\top}\Lambda\vec{v}+\frac{(\alpha+\lambda^2)}{\alpha(1+\lambda)^2}\vec{e}_{-1}^{\top}\Lambda\vec{v}. && (98)
\end{aligned}$$

This shows that $Q_l^{\lambda,p}$ is given by

$$Q_l^{\lambda,p} = -\frac{(\alpha-\lambda)}{\alpha(1+\lambda)^2}\vec{e}_{-1}\vec{e}_{\lambda}^{\top}\Lambda+\frac{(\alpha+\lambda^2)}{\alpha(1+\lambda)^2}\vec{e}_{-1}\vec{e}_{-1}^{\top}\Lambda. \tag{99}$$

Next we compute the matrix elements $[Q_l^{\lambda,p}]_{ij}$ for all $i,j\in\{0,1,\ldots,\alpha\}$. Setting $i=0$, $j=0$ gives

$$\begin{aligned}
[Q_l^{\lambda,p}]_{00} &= -\frac{(\alpha-\lambda)}{\alpha(1+\lambda)^2}(-\lambda)+\frac{(\alpha+\lambda^2)}{\alpha(1+\lambda)^2}(1) && (100)\\
&= \frac{(\alpha-\lambda)\lambda+(\alpha+\lambda^2)}{\alpha(1+\lambda)^2} && (101)\\
&= \frac{\alpha(1+\lambda)}{\alpha(1+\lambda)^2} && (102)\\
&= \frac{1}{1+\lambda} && (103)\\
&= 1-\eta. && (104)
\end{aligned}$$

Setting $i=0$, $j>0$ gives

$$\begin{aligned}
[Q_l^{\lambda,p}]_{0j} &= -\frac{(\alpha-\lambda)}{\alpha(1+\lambda)^2}(-\alpha p_l^{c_j})+\frac{(\alpha+\lambda^2)}{\alpha(1+\lambda)^2}(-\alpha p_l^{c_j}) && (105)\\
&= \frac{\alpha(\alpha-\lambda)-\alpha(\alpha+\lambda^2)}{\alpha(1+\lambda)^2}p_l^{c_j} && (106)\\
&= -\frac{\lambda(1+\lambda)}{(1+\lambda)^2}p_l^{c_j} && (107)\\
&= -\frac{\lambda}{1+\lambda}p_l^{c_j} && (108)\\
&= -\eta p_l^{c_j}. && (109)
\end{aligned}$$

Setting $i>0$, $j=0$ gives

$$\begin{aligned}
[Q_l^{\lambda,p}]_{i0} &= -\frac{(\alpha-\lambda)}{\alpha(1+\lambda)^2}(\lambda)+\frac{(\alpha+\lambda^2)}{\alpha(1+\lambda)^2}(-1) && (110)\\
&= \frac{-\lambda(\alpha-\lambda)-(\alpha+\lambda^2)}{\alpha(1+\lambda)^2} && (111)
\end{aligned}$$

$$= -\frac{\alpha(1+\lambda)}{\alpha(1+\lambda)^2} \tag{112}$$

$$= -\frac{1}{1+\lambda} \tag{113}$$

$$= \eta - 1. \tag{114}$$

Setting $i > 0$, $j > 0$ gives

$$[Q_l^{\lambda,p}]_{ij} = -\frac{(\alpha-\lambda)}{\alpha(1+\lambda)^2}(\alpha p_l^{c_j}) + \frac{(\alpha+\lambda^2)}{\alpha(1+\lambda)^2}(\alpha p_l^{c_j}) \tag{115}$$

$$= \frac{-(\alpha-\lambda)+(\alpha+\lambda^2)}{(1+\lambda)^2} p_l^{c_j} \tag{116}$$

$$= \frac{\lambda(1+\lambda)}{(1+\lambda)^2} p_l^{c_j} \tag{117}$$

$$= \frac{\lambda}{1+\lambda} p_l^{c_j} \tag{118}$$

$$= \eta p_l^{c_j}. \tag{119}$$

We therefore obtain

$$Q_l^{\lambda,p} = \begin{pmatrix} 1-\eta & -p_l^{c_1}\eta & \cdots & -p_l^{c_\alpha}\eta \\ \eta-1 & p_l^{c_1}\eta & \cdots & p_l^{c_\alpha}\eta \\ \vdots & \vdots & \ddots & \vdots \\ \eta-1 & p_l^{c_1}\eta & \cdots & p_l^{c_\alpha}\eta \end{pmatrix}. \tag{120}$$

Finally, plugging this result into $P_l^{\lambda,p} = I - Q_l^{\lambda,p}$ gives

$$P_l^{\lambda,p} = \begin{pmatrix} \eta & p_l^{c_1}\eta & \cdots & p_l^{c_\alpha}\eta \\ 1-\eta & 1-p_l^{c_1}\eta & \cdots & -p_l^{c_\alpha}\eta \\ \vdots & \vdots & \ddots & \vdots \\ 1-\eta & -p_l^{c_1}\eta & \cdots & 1-p_l^{c_\alpha}\eta \end{pmatrix}, \tag{121}$$

which proves the claim. □

**Claim 13.** *$\Theta^{\lambda,p} \equiv \bigotimes_l \Theta_l^{\lambda,p}$ is a valid gauge space for the embedding $\vec{x}_{\text{all}}$.*

*Proof.* This follows from Claim 11 and the fact that $\Theta_l^{\lambda,p}$ is a valid gauge space for $\vec{x}_l$. □

**Claim 14.** *The matrix $P^{\lambda,p}$ that projects $V$ along $G_{\text{all}}$ and into $\Theta^{\lambda,p}$ is an $M \times M$ matrix with elements*

$$P_{s't'}^{\lambda,p} = \prod_{\substack{l \text{ s.t.} \\ s'_l \in \mathcal{A} \\ t'_l \in \mathcal{A}}} \left(\delta_{s'_l t'_l} - p_l^{t'_l}\eta\right) \times \prod_{\substack{l \text{ s.t.} \\ s'_l = * \\ t'_l \in \mathcal{A}}} \left(p_l^{t'_l}\eta\right) \times \prod_{\substack{l \text{ s.t.} \\ s'_l \in \mathcal{A} \\ t'_l = *}} (1-\eta) \times \prod_{\substack{l \text{ s.t.} \\ s'_l = * \\ t'_l = *}} \eta. \tag{122}$$

*Proof.* By Claim 11, $\Theta^{\lambda,p} \equiv \bigotimes_l \Theta_l^{\lambda,p}$ imples that $P^{\lambda,p} \equiv \bigotimes_l P_l^{\lambda,p}$. Eq. 122 follows from expressing the elements of $P_l^{\lambda,p}$ in Eq. 79 as

$$[P^{\lambda,p}]_{cc'} = \begin{cases} \delta_{cc'} - p_l^{c'}\eta & \text{if } c \in \mathcal{A}, c' \in \mathcal{A}, \\ p_l^{c'}\eta & \text{if } c = *, c' \in \mathcal{A}, \\ 1-\eta & \text{if } c \in \mathcal{A}, c' = *, \\ \eta & \text{if } c = *, c' = *, \end{cases} \tag{123}$$

then taking the product across positions $l$ using $c = s'_l$ and $c' = t'_l$. □

**Claim 15.** *If $\lambda$ and $p$ are positive, the $\alpha \times \alpha$ metric*

$$\Lambda_l^{\lambda,p} \equiv \begin{pmatrix} 1 & 0 & \cdots & 0 \\ 0 & \lambda p_l^{c_1} & \cdots & 0 \\ \vdots & \vdots & \ddots & \vdots \\ 0 & 0 & \cdots & \lambda p_l^{c_\alpha} \end{pmatrix} \tag{124}$$

*orthogonalizes $\Theta_l^{\lambda,p}$ and $G_l$.*

*Proof.* Assume that $\Lambda_l^{\lambda,p}$ is diagonal, and thus has the form

$$\Lambda_l^{\lambda,p} = \begin{pmatrix} d_0 & 0 & \cdots & 0 \\ 0 & d_1 & \cdots & 0 \\ \vdots & \vdots & \ddots & \vdots \\ 0 & 0 & \cdots & d_\alpha \end{pmatrix}, \tag{125}$$

for some set of scalars $d_0, \ldots, d_\alpha$. The requirement that $\Lambda_l^{\lambda,p}$ be a metric (and thus positive definite) implies that all $d_0, \ldots, d_\alpha$ must be positive. We now solve for $d_0, \ldots, d_\alpha$ using the matrix equation (Claim 25):

$$\Lambda_l^{\lambda,p} = \left(P_l^{\lambda,p}\right)^\top \Lambda_l^{\lambda,p} \left(P_l^{\lambda,p}\right) + \left(Q_l^{\lambda,p}\right)^\top \Lambda_l^{\lambda,p} \left(Q_l^{\lambda,p}\right), \tag{126}$$

$$\Rightarrow d_i \delta_{ij} = \sum_{k=0}^{\alpha} d_k [P_l^{\lambda,p}]_{ki} [P_l^{\lambda,p}]_{kj} + \sum_{k=0}^{\alpha} d_k [Q_l^{\lambda,p}]_{ki} [Q_l^{\lambda,p}]_{kj}, \tag{127}$$

where $Q_l^{\lambda,p} \equiv I - P_l^{\lambda,p}$. We now solve Eq. 127 for different choices of $i$ and $j$ using the matrix elements of $P_l^{\lambda,p}$ and $Q_l^{\lambda,p}$ computed in the proof of Claim 12. For $i = j = 0$, we get

$$\begin{aligned}
d_0 &= \left[d_0[P_l^{\lambda,p}]_{00}^2 + \sum_{k=1}^{\alpha} d_k [P_l^{\lambda,p}]_{k0}^2\right] + \left[d_0[Q_l^{\lambda,p}]_{00}^2 + \sum_{k=1}^{\alpha} d_k [Q_l^{\lambda,p}]_{k0}^2\right] && (128)\\
&= \left[d_0\eta^2 + \sum_{k=1}^{\alpha} d_k (1-\eta)^2\right] + \left[d_0(1-\eta)^2 + \sum_{k=1}^{\alpha} d_k (\eta-1)^2\right] && (129)\\
&= [d_0\eta^2 + a(1-\eta)^2] + [d_0(1-\eta)^2 + a(\eta-1)^2] && (130)\\
&= d_0[2\eta^2 - 2\eta + 1] + 2a(1-\eta)^2, && (131)\\
\Rightarrow 2a(2-\eta)^2 &= 2d_0\eta(1-\eta), && (132)\\
\Rightarrow a &= d_0 \frac{\eta}{1-\eta} && (133)\\
&= d_0\lambda, && (134)
\end{aligned}$$

where in Eq. 134 we have used $\lambda = \eta/(1-\eta)$. For $i = 0$, $j > 0$, we get

$$\begin{aligned}
0 &= \left[d_0[P_l^{\lambda,p}]_{00}[P_l^{\lambda,p}]_{0j} + \sum_{k=1}^{\alpha} d_k [P_l^{\lambda,p}]_{k0}[P_l^{\lambda,p}]_{kj}\right] + \left[d_0[Q_l^{\lambda,p}]_{00}[Q_l^{\lambda,p}]_{0j} + \sum_{k=1}^{\alpha} d_k [Q_l^{\lambda,p}]_{k0}[Q_l^{\lambda,p}]_{kj}\right] && (135)\\
&= \left[d_0 p_j \eta^2 + \sum_{k=1}^{\alpha} d_k (1-\eta)(\delta_{jk} - p_j\eta)\right] + \left[-d_0 p_j \eta(1-\eta) + \sum_{k=1}^{\alpha} d_k (\eta-1)(p_j\eta)\right] && (136)\\
&= 2d_0 p_j \eta^2 - d_0 p_j \eta - 2a p_j \eta(1-\eta) + d_j(1-\eta) && (137)\\
&= 2(d_0 + a)p_j\eta^2 - (2a + d_0)p_j\eta + d_j(1-\eta) && (138)\\
&= 2d_0(1+\lambda)\frac{\lambda}{1+\lambda} p_j(\eta - 1) + d_0 p_j \eta + d_j(1-\eta), && (139)\\
\Rightarrow d_j &= 2d_0 p_j \lambda + d_0 p_j \frac{\eta}{\eta-1} && (140)\\
&= 2d_0 p_j\lambda - d_0 p_j \lambda && (141)\\
&= d_0 p_j \lambda. && (142)
\end{aligned}$$

The resulting equation,

$$d_i = d_0 p_i \lambda, \quad \text{for all} \quad i = 1, \ldots, \alpha, \tag{143}$$

determines all elements of $\Lambda_l^{\lambda,p}$ up to a multiplicative factor $d_0$. However, we must still verify that Eq. 143 is consistent with the constraints placed on the other elements of $\Lambda_l^{\lambda,p}$ by Eq. 127. For $i > 0$, $j = 0$, we get the same result as for $i = 0$, $j > 0$, because Eq. 127 is symmetric. For $i > 0$, $j > 0$, $i = j$, we get

$$\begin{aligned}
d_i &= \left[d_0[P_l^{\lambda,p}]_{0i}^2 + \sum_{k=1}^{\alpha} d_k [P_l^{\lambda,p}]_{ki}^2\right] + \left[d_0[Q_l^{\lambda,p}]_{0i}^2 + \sum_{k=1}^{\alpha} d_k [Q_l^{\lambda,p}]_{ki}^2\right] && (144)\\
&= \left[d_0 p_i^2 \eta^2 + \sum_{k=1}^{\alpha} d_k (\delta_{ki} - p_i\eta)^2\right] + \left[d_0 p_i^2 \eta^2 + \sum_{k=1}^{\alpha} d_k p_i^2 \eta^2\right] && (145)
\end{aligned}$$

$$
\begin{aligned}
&= \left[d_0 p_i^2\eta^2 + d_i - 2d_i p_i \eta + a p_i^2 \eta^2\right] + \left[d_0 p_i^2 \eta^2 + a p_i^2 \eta^2\right] && (146)\\
&= d_i + 2[(d_0+a)p_i^2\eta^2 - d_i p_i \eta], && (147)\\
\Rightarrow d_i &= (d_0+a)p_i\eta && (148)\\
&= d_0(1+\lambda)\frac{\lambda}{1+\lambda}p_i && (149)\\
&= d_0 p_i \lambda, && (150)
\end{aligned}
$$

which is consistent with Eq. 143. For $i>0$, $j>0$, $i \neq j$, we get

$$
\begin{aligned}
0 &= \left[d_0[P_l^{\lambda,p}]_{0i}[P_l^{\lambda,p}]_{0j} + \sum_{k=1}^{\alpha} d_k [P_l^{\lambda,p}]_{ki}[P_l^{\lambda,p}]_{kj}\right] \\
&\quad + \left[d_0[Q_l^{\lambda,p}]_{0i}[Q_l^{\lambda,p}]_{0j} + \sum_{k=1}^{\alpha} d_k [Q_l^{\lambda,p}]_{ki}[Q_l^{\lambda,p}]_{kj}\right] && (151)\\
&= \left[d_0 p_i p_j \eta^2 + \sum_{k=1}^{\alpha} d_k(\delta_{ki}-p_i\eta)(\delta_{kj}-p_j\eta)\right] + \left[d_0 p_i p_j + \sum_{k=1}^{\alpha} d_k p_i p_j \eta^2\right] && (152)\\
&= 2d_0 p_i p_j \eta^2 + 2a p_i p_j \eta^2 - d_i p_j \eta - p_i d_j \eta && (153)\\
&= 2(d_0+a)p_i p_j \eta^2 - d_i p_j \eta - p_i d_j \eta, && (154)\\
\Rightarrow d_i p_j + p_i d_j &= 2d_0(1+\lambda)\frac{\lambda}{1+\lambda} && (155)\\
&= 2d_0\lambda, && (156)
\end{aligned}
$$

which is also consistent with Eq. 143. We therefore see that Eq. 143 is indeed consistent with Eq. 127. Thus we find that

$$
\Lambda_l^{\lambda,p} = d_0 \begin{pmatrix} 1 & 0 & \cdots & 0 \\ 0 & \lambda p_1 & \cdots & 0 \\ \vdots & \vdots & \ddots & \vdots \\ 0 & 0 & \cdots & \lambda p_\alpha \end{pmatrix}, \tag{157}
$$

orthogonalizes $\Theta_l^{\lambda,p}$ and $G_l$, and is determined up to an unknown positive scalar $d_0$. This proves the claim. □

**Claim 16.** *If $\lambda$ and $p$ are positive, the $M \times M$ metric*

$$
\Lambda_{s't'} \equiv p(s')\lambda^{o(s')}\delta_{s't'} \tag{158}
$$

*orthogonalizes $\Theta^{\lambda,p}$ and $G_{\text{all}}$.*

*Proof.* By Claim 11, $\Theta^{\lambda,p} \equiv \bigotimes_l \Theta_l^{\lambda,p}$ implies that $\Lambda^{\lambda,p} \equiv \bigotimes_l \Lambda_l^{\lambda,p}$. Eq. 158 follows from expressing the elements of $\Lambda_l^{\lambda,p}$ in Eq. 157 as

$$
[\Lambda_l^{\lambda,p}]_{cc'} = \delta_{cc'} p_l^c \times \begin{cases} 1 & \text{if } c = *, \\ \lambda & \text{if } c \in \mathcal{A}, \end{cases} \tag{159}
$$

then taking the product across positions $l$ using $c = s'_l$ and $c' = t'_l$. □

### 6 Trivial gauge, euclidean gauge, and equitable gauge

**Claim 17.** *The trivial gauge $\Theta^{0,p}$ is unaffected by the probability distribution $p$.*

*Proof.* Setting $\lambda = 0$ gives $\eta = 0$, and thus

$$
\begin{aligned}
P_{s't'}^{0,p} &= \prod_{\{l:s'_l\in\mathcal{A} \text{ and } t'_l\in\mathcal{A}\}} \delta_{s'_l t'_l} \times \prod_{\{l:s'_l=*\}} 0 && (160)\\
&= \prod_{\{l:s'_l\in\mathcal{A} \text{ and } t'_l\in\mathcal{A}\}} \delta_{s'_l t'_l} \times \begin{cases} 1 & \text{if } s' \in \mathcal{S} \\ 0 & \text{otherwise} \end{cases} && (161)\\
&= \prod_{\{l:t'_l\in\mathcal{A}\}} \delta_{s'_l t'_l} \times \begin{cases} 1 & \text{if } s' \in \mathcal{S} \\ 0 & \text{otherwise} \end{cases} && (162)
\end{aligned}
$$

$$\Rightarrow P^{0,p}_{s't'} \quad = \quad \begin{cases} 1 & \text{if } s' \in \mathcal{S} \text{ and } x_{t'}(s') = 1 \\ 0 & \text{otherwise} \end{cases} \quad , \tag{163}$$

which does not depend on $p$. $\square$

**Claim 18.** *The euclidean gauge is equal to the embedding space, i.e.,* $\Theta_{\text{eucl}} = S_{\text{all}}$. *Moreover standard* $L_2$ *regularization yields parameters in* $\Theta_{\text{eucl}}$.

*Proof.* In the Euclidean gauge $p(s) = \alpha^{-L}$ for all $s \in \mathcal{S}$. Consequently, $p(s') = \alpha^{-o(s')}$ for all $s' \in \mathcal{S}'$. Because $\lambda = \alpha$ in this gauge as well, the metric $\Lambda$ is given by

$$\Lambda_{s't'} = \delta_{s't'} p(s') \lambda^{o(s')} = \delta_{s't'} \alpha^{-o(s')} \alpha^{o(s')} = \delta_{s't'}. \tag{164}$$

$\Lambda$ is therefore the euclidean metric. Because $\Theta_{\text{eucl}}$ is $\Lambda$-orthogonal to $G_{\text{all}}$,

$$\Theta_{\text{eucl}} = G^{\perp}_{\text{all}} = S_{\text{all}}. \tag{165}$$

And because standard $L_2$ parameter regularization uses a penalty of $\sum_{s'} \theta^2_{s'} = ||\vec{\theta}||^2$, which is the euclidean norm of $\vec{\theta}$, inference using standard $L_2$ regularization yields parameters in the euclidean gauge. This completes the proof. $\square$

**Claim 19.** *In the equitable gauge,* $||\vec{\theta}||^2_{\Lambda} = \sum_{s'} p(s')\theta^2_{s'} = \sum_{s'} \left\langle f^2_{s'} \right\rangle_p$ *where* $f_{s'} \equiv \theta_{s'} x_{s'}$.

*Proof.* The equitable gauge is defined by $\lambda = 1$. Setting $\lambda = 1$ gives $\Lambda_{s't'} = \delta_{s't'} p(s')$, and so

$$\begin{aligned} ||\theta||^2_{\Lambda} &= \sum_{s',t'} \Lambda_{s't'} \theta_{s'} \theta_{t'} && (166) \\ &= \sum_{s'} p(s') \theta^2_{s'} && (167) \\ &= \sum_{s'} \langle x_{s'} \rangle_p \theta^2_{s'} && (168) \\ &= \sum_{s'} \left\langle x^2_{s'} \right\rangle_p \theta^2_{s'} && (169) \\ &= \sum_{s'} \left\langle (\theta_{s'} x_{s'})^2 \right\rangle_p && (170) \\ &= \sum_{s'} \left\langle f^2_{s'} \right\rangle_p && (171) \end{aligned}$$

This completes the proof. $\square$

### 7 Hierarchical gauges

**Definition 18.** *The order* $o(s')$ *of an augmented sequence* $s' \in \mathcal{S}'$ *is defined to be the number of non-star characters in* $s'$, *i.e., the number of positions* $l$ *for which* $s_l \in \mathcal{A}$.

**Definition 19.** *For any two augmented sequences* $s', t' \in \mathcal{S}'$, *we write* $t' \subseteq s'$ *if the sequences matched by* $t'$ *form a subset of those matched by* $s'$. *More formally,* $t' \subseteq s'$ *iff, for all positions* $l$, $s'_l \in \mathcal{A} \Rightarrow t'_l = s'_l$. *Note that* $t' \subseteq s'$ *implies that* $o(t') \geq o(s')$.

**Definition 20.** *For any two augmented sequences* $s', t' \in \mathcal{S}'$, *we write* $t' \succeq s'$ *iff, for all positions* $l$, $s'_l \in \mathcal{A} \Rightarrow t'_l \in \mathcal{A}$, *or equivalently,* $t'_l = * \Rightarrow s'_l = *$. *Note that* $t' \succeq s'$ *implies that* $o(t') \geq o(s')$. *Also note that* $t' \subseteq s' \Rightarrow t' \succeq s'$, *but that the reverse is not true, since* $t' \succeq s'$ *does not require that* $t'_l = s'_l$ *when* $s'_l \in \mathcal{A}$.

For example, if $L = 8$ and $\mathcal{A} = \{\text{A}, \text{C}, \text{G}, \text{T}\}$ is the set of DNA bases, then

$$\begin{aligned} **\text{AT}**\text{C}* \quad &\succeq \quad ***\text{T}**\text{C}*, && (172) \\ **\text{AT}**\text{C}* \quad &\subseteq \quad ***\text{T}**\text{C}*, && (173) \end{aligned}$$

whereas,

$$\begin{aligned} **\text{AT}**\text{G}* \quad &\succeq \quad ***\text{T}**\text{C}*, && (174) \\ **\text{AT}**\text{G}* \quad &\not\subseteq \quad ***\text{T}**\text{C}*\,. && (175) \end{aligned}$$

**Definition 21.** *A hierarchical model is a an all-order interaction model in which $\theta_{s'} = 0$ for all augmented sequences $s'$ in a set $\mathcal{Z}' \subseteq \mathcal{S}'$ (the "zero set" of the model) that has the following property: for every $s' \in \mathcal{Z}'$ and $t' \in \mathcal{S}'$, $t' \succeq s'$ implies that $t' \in \mathcal{Z}'$.*

**Claim 20.** *Hierarchical gauges preserve the form of hierarchical models. Specifically, if $\vec{\theta}$ are the parameters of a hierarchical model having zero set $\mathcal{Z}'$, then $\vec{\theta}_{\text{fixed}} = P^{\infty,p}\vec{\theta}$ are also parameters of a hierarchical model with zero set $\mathcal{Z}'$.*

*Proof.* Setting $\lambda = \infty$ in Eq. 22 of the main text, we see that the elements of $P^{\infty;p}_{s't'}$ can written

$$P^{\infty;p}_{s't'} = \prod_{\substack{l \text{ s.t.} \\ s'_l \in \mathcal{A} \\ t'_l \in \mathcal{A}}} \left(\delta_{s'_l t'_l} - p_l^{t'_l}\right) \times \prod_{\substack{l \text{ s.t.} \\ s'_l = * \\ t'_l \in \mathcal{A}}} p_l^{t'_l} \times \prod_{\substack{l \text{ s.t.} \\ s'_l \in \mathcal{A} \\ t'_l = *}} 0, \tag{176}$$

$$= \prod_{\substack{l \text{ s.t.} \\ s'_l \in \mathcal{A} \\ t'_l \in \mathcal{A}}} \left(\delta_{s'_l t'_l} - p_l^{t'_l}\right) \times \prod_{\substack{l \text{ s.t.} \\ s'_l = * \\ t'_l \in \mathcal{A}}} p_l^{t'_l} \times \begin{cases} 1 & \text{if } t' \succeq s' \\ 0 & \text{otherwise} \end{cases} \tag{177}$$

$$= \prod_{\{l : s'_l \in \mathcal{A}\}} \left(\delta_{s'_l t'_l} - p_l^{t'_l}\right) \times \prod_{\{l : s'_l = *\}} p_l^{t'_l} \times \begin{cases} 1 & \text{if } t' \succeq s' \\ 0 & \text{otherwise} \end{cases} . \tag{178}$$

Here we used Def. 20 in going from Eq. 176 to Eq. 177, and we used the fact that $t' \succeq s'$ implies that $s'_l \in \mathcal{A} \Rightarrow t'_l \in \mathcal{A}$, as well as the fact that $t'_l = * \Rightarrow p_l^{t'_l} = 1$, in going from Eq. 177 to Eq. 178. Now assume $\vec{\theta}$ are the parameters of a hierarchical model with zero set $\mathcal{Z}'$, and choose any $s' \in \mathcal{Z}'$. Then in the hierarchical gauge $\Theta^{\infty,p}_{s't'}$,

$$\theta^{\text{fixed}}_{s'} = \sum_{t' \in \mathcal{S}'} P^{\infty;p}_{s't'} \theta_{t'}. \tag{179}$$

$$= \sum_{t' \succeq s'} \theta_{t'} \prod_{\{l : s'_l \in \mathcal{A}\}} \left(\delta_{s'_l t'_l} - p_l^{t'_l}\right) \prod_{\{l : s'_l = *\}} p_l^{t'_l} \tag{180}$$

$$= 0 \tag{181}$$

because $t' \succeq s'$ implies that $t' \in \mathcal{Z}'$ and thus that $\theta_{t'} = 0$ for all $t'$ in the sum. We conclude that $\theta^{\text{fixed}}_{s'} = 0$ for every $s' \in \mathcal{Z}'$, i.e., $\vec{\theta}_{\text{fixed}}$ are the parameters of a hierarchical model defined by zero set $\mathcal{Z}'$. □

**Claim 21.** *Parameters $\vec{\theta}_{\text{fixed}}$ in the hierarchical gauge $\Theta^{\infty,p}$ satisfy the marginalization constraint*

$$\sum_{c_k} p_{l_k}^{c_k} \theta^{c_1 \ldots c_K}_{l_1 \ldots l_K, \text{fixed}} = 0 \tag{182}$$

*for every $K = 1, \ldots, L$, every subset of positions $\{l_1, \ldots, l_K\}$, and every choice of $k = 1, \ldots, K$.*

*Proof.* From Eq. 180 in the proof of Claim 20, parameters $\vec{\theta}$ in the hierarchical gauge $\Theta^{\infty,p}$ are given in terms of unfixed parameters $\vec{\theta}$ via

$$\theta^{\text{fixed}}_{s'} = \sum_{t' \in \mathcal{S}'} P^{\infty;p}_{s't'} \theta_{t'} = \sum_{t' \succeq s'} \theta_{t'} \prod_{\{l : s'_l \in \mathcal{A}\}} \left(\delta_{s'_l t'_l} - p_l^{t'_l}\right) \prod_{\{l : s'_l = *\}} p_l^{t'_l}. \tag{183}$$

Now choose any $K \in \{1, \ldots, L\}$, any set of positions $\sigma = \{l_1, \ldots, l_K\}$, and any index $k \in \{1, \ldots, K\}$. Define $u' \in \mathcal{S}'$ to be the augmented sequence for which $u'_{l_i} = c_i$ for all $i = 1, \ldots, K$, and that has $u'_l = *$ for all $l \notin \sigma$. Further define $\mathcal{S}_{u',k} \subseteq \mathcal{S}'$ to bet the set of augmented sequences obtained by replacing the character at position $k$ in $u'$ with the $\alpha$ different characters in $\mathcal{A}$. To reduce the notational burden, we use $i$ as a synonym for $l_i$ when $i = 1, \ldots, K$, and use $i = K+1, \ldots, L$ to denote positions not in $\sigma$. We find that

$$\sum_{c_k} p_k^{c_k} \theta^{c_1 \ldots c_K}_{1 \ldots K, \text{fixed}} = \sum_{s' \in \mathcal{S}_{u',k}} p_k^{s'_k} \theta_{s'} \tag{184}$$

$$= \sum_{s' \in \mathcal{S}_{u',k}} p_k^{s'_k} \sum_{t' \subseteq s'} \theta_{t'} \prod_{i=1}^{K} \left(\delta_{s'_i t'_i} - p_i^{t'_i}\right) \prod_{i=K+1}^{L} p_i^{t'_i} \tag{185}$$

$$= \sum_{t' \subseteq u'} \sum_{s' \in \mathcal{S}_{u',k}} \theta_{t'} p_k^{s'_k} \prod_{i=1}^{K} \left(\delta_{s'_i t'_i} - p_i^{t'_i}\right) \prod_{i=K+1}^{L} p_i^{t'_i} \tag{186}$$

$$= \sum_{t' \subseteq u'} \sum_{c \in \mathcal{A}} \theta_{t'} p_k^c (\delta_{ct'_k} - p_k^{t'_k}) \prod_{\substack{i=1 \\ i \neq k}}^{K} \Big(\delta_{u'_i t'_i} - p_i^{t'_i}\Big) \prod_{i=K+1}^{L} p_i^{t'_i} \tag{187}$$

$$= \sum_{t' \subseteq u'} \theta_{t'} \left[\sum_{c \in \mathcal{A}} p_k^c (\delta_{ct'_k} - p_k^{t'_k})\right] \prod_{\substack{i=1 \\ i \neq k}}^{K} \Big(\delta_{u'_i t'_i} - p_i^{t'_i}\Big) \prod_{i=K+1}^{L} p_i^{t'_i} \tag{188}$$

$$= 0. \tag{189}$$

In going from Eq. 184 to Eq. 185 we used Eq. 183. In going from Eq. 185 to Eq. 186 we used the fact that $t' \subseteq s'$ and $t' \subseteq u'$ are the same condition on $t'$ when $s' \in \mathcal{S}_{u',k}$. In going from Eq. 186 to Eq. 187, we eliminated $s'$ by separating the case $i = k$ out of the product over $i = 1, \ldots, K$, by replacing $s'_k \to c$, and by replacing $s'_i \to u'_i$ for all $i \neq k$. In going from Eq. 187 to Eq. 188, we collected in brackets all quantities that depend on $c$. And in going from Eq. 188 to Eq. 189, we use the fact that the term in brackets vanishes:

$$\sum_{c \in \mathcal{A}} p_k^c \left(\delta_{ct'_k} - p_k^{t'_k}\right) = \left(\sum_{c \in \mathcal{A}} p_k^c \delta_{ct'_k}\right) - \left(\sum_{c \in \mathcal{A}} p_k^c\right) p_k^{t'_k} = p_k^{t'_k} - p_k^{t'_k} = 0. \tag{190}$$

This proves the claim. □

**Definition 22.** *The $\mathcal{A}$-positions of an augmented sequence $s'$ are the positions $l$ such that $s'_l \in \mathcal{A}$. Similarly, the $*$-positions of $s'$ are the positions $l$ such that $s'_l = *$.*

**Definition 23.** *An augmented sequence orbit $\sigma$ is a set comprising all augmented sequences that have a specified set of $\mathcal{A}$-positions (or equivalently, a specified set of $*$-positions). The order of the orbit, $o(\sigma)$, is defined to be the order of all $s' \in \sigma$. The term "orbit" comes from the fact that such sets are formed from the orbit of $s'$ under the group of position-specific character permutations; see ref. [1].*

**Claim 22.** *Let $f(s) = \sum_{t'} \theta_{t'} x_{t'}(s)$ be an activity landscape and $p$ be a positive probability distribution. Define the expectation value of $f$ with respect to $p$ conditioned on $s' \in \mathcal{S}'$ to be $\langle f|s' \rangle_p = \frac{1}{p(s')} \sum_{s \in s'} p(s) f(s)$. Then when $\vec{\theta}$ is in the hierarchical gauge,*

$$\langle f|s' \rangle_p = \sum_{t' \supseteq s'} \theta_{t'}. \tag{191}$$

*This claim is readily extended to non-positive probability distributions $p$ by defining $\langle f|s' \rangle_p = \lim_{\epsilon \to 0^+} \langle f|s' \rangle_{p_\epsilon}$, where $p_\epsilon$ is a regularized version of $p$ given by*

$$p_\epsilon(s) = \prod_l \left[(1-\epsilon) p_l^{s_l} + \frac{\epsilon}{\alpha}\right], \tag{192}$$

*with $p_l$ being the position-specific factors of $p$.*

*Proof.* Assume that $p$ is positive, and that $\vec{\theta}$ is in the hierarchical gauge. Then,

$$\langle f|s' \rangle_p = \frac{1}{p(s')} \sum_{s \in s'} p(s) \sum_{t'} \theta_{t'} x_{t'}(s) \tag{193}$$

$$= \frac{1}{p(s')} \sum_{t'} \theta_{t'} \sum_{s \in s'} p(s) x_{t'}(s) \tag{194}$$

$$= \frac{1}{p(s')} \sum_{t'} \theta_{t'} \sum_{s \in s' \cap t'} p(s) \tag{195}$$

$$= \frac{1}{p(s')} \sum_{t'} p(s' \cap t') \theta_{t'} \tag{196}$$

$$= \frac{1}{p(s')} \sum_{\tau} \sum_{t' \in \tau} p(s' \cap t') \theta_{t'}. \tag{197}$$

where $\sum_\tau$ denotes a sum over all augmented sequence orbits $\tau$. Now let $l_1, \ldots, l_K, m_1, \ldots, m_J$ denote the $\mathcal{A}$-positions of $s'$, let $l_1, \ldots, l_K, n_1, \ldots, n_I$ denote the $\mathcal{A}$-positions of $\tau$, and assume $l_1, \ldots, l_K, m_1, \ldots, m_J, n_1, \ldots, n_I$ are distinct. Then for each orbit $\tau$,

$$\sum_{t' \in \tau} p(s' \cap t') \theta_{t'} = \sum_{t'_{l_1} \cdots t'_{l_K}} \sum_{t'_{n_1} \cdots t'_{n_I}} p_{l_1}^{s'_{l_1}} \cdots p_{l_K}^{s'_{l_K}} p_{m_1}^{s'_{m_1}} \cdots p_{m_J}^{s'_{m_J}} p_{n_1}^{t'_{n_1}} \cdots p_{n_I}^{t'_{n_I}} \delta_{s'_{l_1} t'_{l_1}} \cdots \delta_{s'_{l_K} t'_{l_K}} \theta_{l_1 \cdots l_K n_1 \cdots n_I}^{t'_{l_1} \cdots t'_{l_K} t'_{n_1} \cdots t'_{n_I}} \tag{198}$$

$$= p_{l_1}^{s'_{l_1}} \cdots p_{l_K}^{s'_{l_K}} p_{m_1}^{s'_{m_1}} \cdots p_{m_J}^{s'_{m_J}} \sum_{t'_{l_1}\cdots t'_{l_K}} \delta_{s'_{l_1}t'_{l_1}} \cdots \delta_{s'_{l_K}t'_{l_K}} \sum_{t'_{n_1}\cdots t'_{n_I}} p_{n_1}^{t'_{n_1}} \cdots p_{n_I}^{t'_{n_I}} \theta_{l_1\cdots l_K n_1\cdots n_I}^{t'_{l_1}\cdots t'_{l_K} t'_{n_1}\cdots t'_{n_I}}. \quad (199)$$

Noting that

$$\delta_{s'_{l_1}t'_{l_1}} \cdots \delta_{s'_{l_K}t'_{l_K}} = \begin{cases} 1 & \text{if } s'_l = t'_l \text{ for all } l = l_1, \ldots, l_K, \\ 0 & \text{otherwise}, \end{cases} \quad (200)$$

and that by Claim 21,

$$\sum_{t'_{n_1}\cdots t'_{n_I}} p_{n_1}^{t'_{n_1}} \cdots p_{n_I}^{t'_{n_I}} \theta_{l_1\cdots l_K n_1\cdots n_I}^{t'_{l_1}\cdots t'_{l_K} t'_{n_1}\cdots t'_{n_I}} = \begin{cases} \theta_{l_1\cdots l_K}^{t'_{l_1}\cdots t'_{l_K}} & \text{if } I = 0, \\ 0 & \text{otherwise}, \end{cases} \quad (201)$$

we see that,

$$\sum_{t'\in\tau} p(s'\cap t')\theta_{t'} = p_{l_1}^{s'_{l_1}} \cdots p_{l_K}^{s'_{l_K}} p_{m_1}^{s'_{m_1}} \cdots p_{m_J}^{s'_{m_J}} \sum_{t'_{l_1}\cdots t'_{l_K}} \theta_{l_1\cdots l_K}^{t'_{l_1}\cdots t'_{l_K}} \times \begin{cases} 1 \text{ if } t'_l \in \mathcal{A} \Rightarrow s'_l = t'_l, \text{ for all } l = l_1, \ldots, l_K, \\ 0 \text{ otherwise.} \end{cases} \quad (202)$$

$$= p(s') \sum_{t'\in\tau} \theta_{t'} \begin{cases} 1 \text{ if } s' \subseteq t', \\ 0 \text{ otherwise.} \end{cases} \quad (203)$$

Consequently,

$$\langle f|s'\rangle_p = \frac{1}{p(s')} \sum_{\tau} p(s') \sum_{t'\in\tau} \theta_{t'} \begin{cases} 1 \text{ if } s' \subseteq t', \\ 0 \text{ otherwise}, \end{cases} \quad (204)$$

$$= \sum_{t'} \theta_{t'} \begin{cases} 1 \text{ if } s' \subseteq t', \\ 0 \text{ otherwise}, \end{cases} \quad (205)$$

$$= \sum_{t'\supseteq s'} \theta_{t'}. \quad (206)$$

This completes the proof for the case where $p$ is positive. The proof for non-positive $p$ follows from the definition and the continuity of projection matrix elements $P_{s't'}^{\infty,p}$ (Eq. 176) with respect to $p$ and the definition $\langle f|s'\rangle_p = \lim_{\epsilon\to 0^+} \langle f|s'\rangle_{p_\epsilon}$. □

**Definition 24.** *The orbital component of an activity landscape $f = \sum_{s'} \theta_{s'} x_{s'}$ corresponding to an augmented sequence orbit $\sigma$ is defined to be $f_\sigma(s) = \sum_{s'\in\sigma} \theta_{s'} x_{s'}(s)$.*

**Claim 23.** *Given an activity landscape $f$, and augmented sequence orbits $\sigma$ and $\tau$,*

$$\langle f_\sigma\rangle_p = \begin{cases} \theta_0 & \text{if } o(\sigma) = 0, \\ 0 & \text{otherwise}, \end{cases} \quad (207)$$

*and $\langle f_\sigma f_\tau\rangle_p = \delta_{\sigma\tau} \langle f_\sigma^2\rangle_p$ where*

$$\langle f_\sigma^2\rangle_p = \sum_{s'\in\sigma} p(s')\theta_{s'}^2. \quad (208)$$

*Proof.* Eq. 207 follows directly from Claim 21:

$$\langle f_\sigma\rangle_p = \sum_{s'\in\sigma} p(s')\theta_{s'} \begin{cases} \theta_0 & \text{if } o(\sigma) = 0, \\ 0 & \text{otherwise}, \end{cases} \quad (209)$$

Let $l_1, \ldots, l_K, m_1, \ldots, m_J$ denote the $\mathcal{A}$-positions of $\sigma$, let $l_1, \ldots, l_K, n_1, \ldots, n_I$ denote the $\mathcal{A}$-positions of $\tau$, and assume $l_1, \ldots, l_K, m_1, \ldots, m_J, n_1, \ldots, n_I$ are distinct. Then,

$$\langle f_\sigma f_\tau\rangle_p = \sum_u p(u) \left[\sum_{s'\in\sigma} \theta_{s'} x_{s'}(u)\right] \left[\sum_{t'\in\tau} \theta_{t'} x_{t'}(u)\right] \quad (210)$$

$$= \sum_{s'\in\sigma} \sum_{t'\in\tau} \theta_{s'}\theta_{t'} \sum_u p(u) x_{s'\cap t'}(u) \quad (211)$$

$$= \sum_{s'\in\sigma} \sum_{t'\in\tau} p(s'\cap t')\theta_{s'}\theta_{t'} \quad (212)$$

$$= \sum_{s'_{l_1}\cdots s'_{l_K}} \sum_{s'_{m_1}\cdots s'_{m_J}} \sum_{t'_{l_1}\cdots t'_{l_K}} \sum_{t'_{n_1}\cdots t'_{n_I}} p_{l_1}^{s'_{l_1}} \cdots p_{l_K}^{s'_{l_K}} p_{m_1}^{s'_{m_1}} \cdots p_{m_J}^{s'_{m_J}} p_{n_1}^{t'_{n_1}} \cdots p_{n_I}^{t'_{n_I}} \times \quad (213)$$

$$\delta_{s'_{l_1}t'_{l_1}}\cdots\delta_{s'_{l_K}t'_{l_K}}\theta^{s'_{l_1}\cdots s'_{l_K}s'_{m_1}\cdots s'_{m_J}}_{l_1\cdots l_K m_1\cdots m_J}\theta^{t'_{l_1}\cdots t'_{l_K}t'_{n_1}\cdots t'_{n_I}}_{l_1\cdots l_K n_1\cdots n_I} \tag{214}$$

$$= \sum_{s'_{l_1}\cdots s'_{l_K}}\sum_{t'_{l_1}\cdots t'_{l_K}} p^{s'_{l_1}}_{l_1}\cdots p^{s'_{l_K}}_{l_K}\delta_{s'_{l_1}t'_{l_1}}\cdots\delta_{s'_{l_K}t'_{l_K}}\times \tag{215}$$

$$\left[\sum_{s'_{m_1}\cdots s'_{m_J}} p^{s'_{m_1}}_{m_1}\cdots p^{s'_{m_J}}_{m_J}\theta^{s'_{l_1}\cdots s'_{l_K}s'_{m_1}\cdots s'_{m_J}}_{l_1\cdots l_K m_1\cdots m_J}\right]\times\left[\sum_{t'_{n_1}\cdots t'_{n_I}} p^{t'_{n_1}}_{n_1}\cdots p^{t'_{n_I}}_{n_I}\theta^{t'_{l_1}\cdots t'_{l_K}t'_{n_1}\cdots t'_{n_I}}_{l_1\cdots l_K n_1\cdots n_I}\right]. \tag{216}$$

Because

$$\sum_{s'_{m_1}\cdots s'_{m_J}} p^{s'_{m_1}}_{m_1}\cdots p^{s'_{m_J}}_{m_J}\theta^{s'_{l_1}\cdots s'_{l_K}s'_{m_1}\cdots s'_{m_J}}_{l_1\cdots l_K m_1\cdots m_J} = \begin{cases}\theta^{s'_{l_1}\cdots s'_{l_K}}_{s_1\cdots s_K} & \text{if } J=0,\\ 0 & \text{otherwise,}\end{cases} \tag{217}$$

and

$$\sum_{t'_{n_1}\cdots t'_{n_I}} p^{t'_{n_1}}_{n_1}\cdots p^{t'_{n_I}}_{n_I}\theta^{t'_{l_1}\cdots t'_{l_K}t'_{n_1}\cdots t'_{n_I}}_{l_1\cdots l_K n_1\cdots n_I} = \begin{cases}\theta^{t'_{l_1}\cdots t'_{l_K}}_{l_1\cdots l_K} & \text{if } I=0,\\ 0 & \text{otherwise,}\end{cases} \tag{218}$$

we see that the summand in Eq. 215 vanishes unless $J = 0$ and $I = 0$. This is equivalent to the requirement that $\sigma = \tau$. Therefore,

$$\langle f_\sigma f_\tau\rangle_p = \delta_{\sigma\tau}\sum_{s'_{l_1}\cdots s'_{l_K}}\sum_{t'_{l_1}\cdots t'_{l_K}} p^{s'_{l_1}}_{l_1}\cdots p^{s'_{l_K}}_{l_K}\delta_{s'_{l_1}t'_{l_1}}\cdots\delta_{s'_{l_K}t'_{l_K}}\theta^{s'_{l_1}\cdots s'_{l_K}}_{s_1\cdots s_K}\theta^{t'_{l_1}\cdots t'_{l_K}}_{l_1\cdots l_K} \tag{219}$$

$$= \delta_{\sigma\tau}\sum_{s'\in\sigma}\sum_{t'\in\tau} p(s')\delta_{s't'}\theta_{s'}\theta_{t'} \tag{220}$$

$$= \delta_{\sigma\tau}\sum_{s'\in\sigma} p(s')\theta^2_{s'} \tag{221}$$

$$= \delta_{\sigma\tau}\left\langle f^2_\sigma\right\rangle_p \tag{222}$$

where

$$\left\langle f^2_\sigma\right\rangle_p = \sum_{s'\in\sigma} p(s')\theta^2_{s'}. \tag{223}$$

□

**Definition 25.** *Given $k \in \{0, 1, \ldots, L\}$, the k'the order component of an activity landscape $f = \sum_{s'}\theta_{s'}x_{s'}$ is defined to be*

$$f_k(s) = \sum_{s':o(s')=k}\theta_{s'}x_{s'}(s). \tag{224}$$

**Claim 24.** *Given an activity landscape $f = \sum_{s'}\theta_{s'}x_{s'}$, and parameters $\vec{\theta}$ expressed in the hierarchical gauge,*

$$\mathrm{var}_p[f] = \sum_{k=0}^{L}\mathrm{var}_p[f_k], \quad \textit{where} \quad \mathrm{var}_p[f_k] = \begin{cases}\sum_{s':o(s')=k}p(s')\theta^2_{s'} & \textit{if } k\geq 1,\\ 0 & \textit{if } k=0.\end{cases} \tag{225}$$

*Proof.* First we decompose each $f_k$ into a sum of $f_\sigma$ over augmented sequence orbits $\sigma$ of order $k$:

$$f_k = \sum_{\sigma:o(\sigma)=k} f_\sigma. \tag{226}$$

next, by Claim 23,

$$\langle f_k\rangle_p = \sum_{\sigma:o(\sigma)=k}\langle f_\sigma\rangle_p, \tag{227}$$

and

$$\left\langle f^2_k\right\rangle_p = \left\langle\sum_{\sigma:o(\sigma)=k} f_\sigma\sum_{\tau:o(\tau)=k} f_\tau\right\rangle_p \tag{228}$$

$$= \sum_{\sigma:o(\sigma)=k}\sum_{\tau:o(\tau)=k}\langle f_\sigma f_\tau\rangle_p \tag{229}$$

$$= \sum_{\sigma:o(\sigma)=k} \sum_{\tau:o(\tau)=k} \delta_{\sigma\tau} \left\langle f_\sigma^2 \right\rangle \tag{230}$$

$$= \sum_{\sigma:o(\sigma)=k} \left\langle f_\sigma^2 \right\rangle \tag{231}$$

$$= \sum_{\sigma:o(\sigma)=k} \sum_{s'\in\sigma} p(s')\theta_{s'}^2 \tag{232}$$

$$= \sum_{s':o(s')=k} p(s')\theta_{s'}^2. \tag{233}$$

Consequently

$$\begin{aligned}
\mathrm{var}_p[f_k] &= \left\langle f_k^2 \right\rangle_p - \left\langle f_k \right\rangle_p^2 && (234)\\
&= \sum_{s':o(s')=k} p(s')\theta_{s'}^2 - \delta_{k0}\theta_0^2 && (235)\\
&= \begin{cases} \sum_{s':o(s')=k} p(s')\theta_{s'}^2 & \text{if} \quad k \geq 1, \\ 0 & \text{if} \quad k = 0. \end{cases} && (236)
\end{aligned}$$

Finally,

$$\begin{aligned}
\mathrm{var}_p[f] &= \left\langle f^2 \right\rangle_p - \left\langle f \right\rangle_p^2 && (237)\\
&= \left\langle \sum_\sigma f_\sigma \sum_\tau f_\tau \right\rangle_p - \left\langle \sum_\sigma f_\sigma \right\rangle_p^2 && (238)\\
&= \sum_\sigma \sum_\tau \left\langle f_\sigma f_\tau \right\rangle - \sum_\sigma \left\langle f_\sigma \right\rangle_p^2 && (239)\\
&= \sum_\sigma \left[ \left\langle f_\sigma^2 \right\rangle_p - \left\langle f_\sigma \right\rangle^2 \right] && (240)\\
&= \sum_{k=0}^{L} \sum_{\sigma:o(\sigma)=k} \left[ \left\langle f_\sigma^2 \right\rangle_p - \left\langle f_\sigma \right\rangle^2 \right] && (241)\\
&= \sum_{k=0}^{L} \mathrm{var}_p[f_k]. && (242)
\end{aligned}$$

This completes the proof. □

### 8 Hierarchical gauges of an empirical landscape for protein GB1

**Gauge-fixing formula for the all-order interaction model** Using the formula for $P^{\infty,p}$ in Eq. 176 to compute

$$\vec{\theta}_{\text{fixed}} = P^{\infty,p}\vec{\theta} \tag{243}$$

for an all-order interaction model in which only the zero-order, first-order, and second-order parameters are nonzero, one finds that

$$\begin{aligned}
\theta_{0,\text{fixed}} &= \theta_0 + \sum_l \sum_c p_l^c \theta_l^c + \sum_l \sum_{l'>l} \sum_{c,c'} p_l^c p_{l'}^{c'} \theta_{ll'}^{cc'}, && (244)\\
\theta_{l,\text{fixed}}^c &= \sum_{c'} (\delta_{cc'} - p_l^{c'})\theta_l^c + \sum_{l'<l} \sum_{c',c''} (\delta_{cc'} - p_l^{c'}) p_{l'}^{c''} \theta_{ll'}^{cc'} + \sum_{l'>l} \sum_{c',c''} (\delta_{cc'} - p_l^{c'}) p_{l'}^{c''} \theta_{l'l}^{c'c}, && (245)\\
\theta_{ll',\text{fixed}}^{cc'} &= \sum_{c'',c'''} (\delta_{cc''} - p_l^{c''})(\delta_{c'c'''} - p_{l'}^{c'''})\theta_{ll'}^{c''c'''}, && (246)\\
\theta_{l_1 \ldots l_K,\text{fixed}}^{c_1 \ldots c_K} &= 0 \quad \text{for all} \quad K = 3, \ldots, L. && (247)
\end{aligned}$$

Ignoring the formula for parameters of order three or greater, one thus obtains the gauge-fixing formulae for the parameters of the pairwise-interaction model. These are the formulas used for the computations in Fig. 4 and Fig. 5 of the main text. The specific choices for $p$ used in these figures are given below.

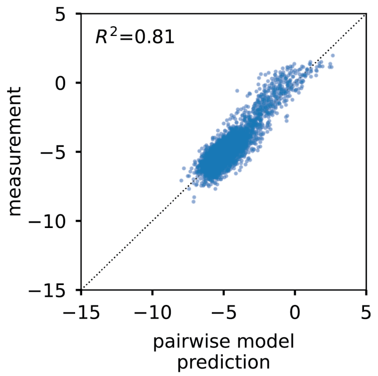


Figure S1: Performance of the pairwise-interaction model for protein GB1. Axes reflect $\log_2$ enrichment ratios. Each dot represents a randomly chosen variant GB1 protein assayed by [2]. For clarity, only 5,000 of the ∼160,000 assayed GB1 variants are shown.

**Region-specific distributions.** In Fig. 4D, the probability distributions $p(s) = \prod_{l=1}^{4} p_l^{s_l}$ for the four different regions (global, region 1, region 2, region 3) were defined as follows.

- Uniform: For $l \in \{1, 2, 3, 4\}$, $p_l^c = \frac{1}{20}$.
- Region 1: For $l \in \{1, 2, 4\}$, $p_l^c = \frac{1}{20}$; for $l = 3$, $p_l^c = \delta_{c\mathrm{G}}$.
- Region 2: For $l \in \{1, 2\}$, $p_l^c = \frac{1}{20}$; for $l = 3$, $p_l^c = \frac{1}{2}\delta_{c\mathrm{L}} + \frac{1}{2}\delta_{c\mathrm{F}}$; for $l = 4$, $p_l^c = \delta_{c\mathrm{G}}$.
- Region 3: For $l \in \{1, 2\}$, $p_l^c = \frac{1}{20}$; for $l = 3$, $p_l^c = \frac{1}{2}\delta_{c\mathrm{C}} + \frac{1}{2}\delta_{c\mathrm{A}}$; for $l = 4$, $p_l^c = \delta_{c\mathrm{A}}$.

Here $l = 1, 2, 3, 4$ are used to denote protein positions $39, 40, 41, 54$, respectively, and $c$ indexes all $\alpha = 20$ possible amino acids in $\mathcal{A}$.

### 9 Appendix

**Claim 25.** *Let $V_1$ and $V_2$ be two subspaces of a vector space $V$ such that any vector in $V$ can be uniquely decomposed into the sum of a vector in $V_1$ and a vector in $V_2$. Let $P_1$ be the projection into $V_1$ along $V_2$, and $P_2$ be the projection into $V_2$ along $V_1$. Let $\Lambda$ be a symmetric positive definite matrix acting on $V$. Then the following three statements are equivalent.*

1. *$V_1$ and $V_2$ are $\Lambda$-orthogonal, i.e., $\vec{v}_1^\top \Lambda \vec{v}_2 = 0$ for all $\vec{v}_1 \in V_1$ and $\vec{v}_2 \in V_2$.*
2. *For any fixed $\vec{v}_1 \in V_1$, $\mathrm{argmin}_{\vec{v}_2 \in V_2} (\vec{v}_1 + \vec{v}_2)^\top \Lambda (\vec{v}_1 + \vec{v}_2) = \vec{0}$.*
3. *$\Lambda = P_1^\top \Lambda P_1 + P_2^\top \Lambda P_2$.*

*Proof.* We prove equivalence of the three statements (denoted 1, 2, and 3) as follows.

- $1 \Rightarrow 2$: Assume that 1 is true. Then for all $\vec{v}_1 \in V_1$ and $\vec{v}_2 \in V_2$, $(\vec{v}_1 + \vec{v}_2)^\top \Lambda (\vec{v}_1 + \vec{v}_2) = \vec{v}_1^\top \Lambda \vec{v}_1 + \vec{v}_2^\top \Lambda \vec{v}_2 \geq \vec{v}_1^\top \Lambda \vec{v}_1$. Because equality obtains only when $\vec{v}_2 = \vec{0}$, $\vec{0}$ is the unique $\vec{v}_2 \in V$ that minimizes $(\vec{v}_1 + \vec{v}_2)^\top \Lambda (\vec{v}_1 + \vec{v}_2)$. This proves 2, thereby establishing that $1 \Rightarrow 2$.
- $2 \Rightarrow 1$: Assume that 1 is not true, i.e., there exists $\vec{v}_1 \in V_1$ and $\vec{v}_2 \in V_2$ such that $\vec{v}_1^\top \Lambda \vec{v}_2 \neq 0$. Then $\frac{d}{d\epsilon}(\vec{v}_1 + \epsilon \vec{v}_2)^\top \Lambda (\vec{v}_1 + \epsilon \vec{v}_2) = 2\vec{v}_1^\top \Lambda \vec{v}_2 + 2\epsilon \vec{v}_2^\top \Lambda \vec{v}_2$ is nonzero at $\epsilon = 0$. This contradicts 2, thereby establishing that $\neg 1 \Rightarrow \neg 2$ and hence $2 \Rightarrow 1$.
- $1 \Rightarrow 3$: Assume that 1 is true, and choose any $\vec{v}, \vec{w} \in V$. Then $\vec{v}^\top \Lambda \vec{w} = (P_1 \vec{v} + P_2 \vec{v})^\top \Lambda (P_1 \vec{w} + P_2 \vec{w}) = (P_1 \vec{v})^\top \Lambda (P_1 \vec{w}) + (P_2 \vec{v})^\top \Lambda (P_2 \vec{w}) = \vec{v}^\top \left[ P_1^\top \Lambda P_1 + P_2^\top \Lambda P_2 \right] \vec{w}$. This proves 3, thereby establishing that $1 \Rightarrow 3$.
- $3 \Rightarrow 1$: Assume that 3 is true. Then given any $\vec{v}_1 \in V_1$ and $\vec{v}_2 \in V_2$, $\vec{v_1}^\top \Lambda \vec{v}_2 = (P_1 \vec{v_1})^\top \Lambda (P_1 \vec{v}_2) + (P_2 \vec{v_1})^\top \Lambda (P_2 \vec{v}_2) = 0$ since $P_1 \vec{v}_2 = P_2 \vec{v_1} = \vec{0}$. This proves 1, thereby establishing that $3 \Rightarrow 1$.

□
